# Partial prion cross-seeding between fungal and mammalian amyloid signaling motifs

**DOI:** 10.1101/2020.07.22.215483

**Authors:** Thierry Bardin, Asen Daskalov, Sophie Barrouilhet, Alexandra Granger-Farbos, Bénédicte Salin, Corinne Blancard, Sven J. Saupe, Virginie Coustou

**Affiliations:** Non-Self Recognition in Fungi, Institut de Biochimie et de Génétique Cellulaire (CNRS UMR 5095, Université de Bordeaux), 33077 Bordeaux, France

**Author notes:** Current Address: Universite de Pau et des Pays de l’Adour, E2S UPPA, CNRS, IPREM UMR 5254, Hélioparc, 64053 Pau, France. Corresponding authors (VC); (SJS).

**Keywords:** amyloid, prion, programmed cell death, RHIM, RIP1, RIP3

## Abstract

In filamentous fungi, NLR-based signalosomes activate downstream membrane-targeting cell-death inducing proteins by a mechanism of amyloid templating. In the species *Podospora anserina*, two such signalosomes, NWD2/HET-S and FNT1/HELLF have been described. An analogous system, involving a distinct amyloid signaling motif termed PP was also identified in the genome of the species *Chaetomium globosum* and studied using heterologous expression in *Podospora anserina*. The PP-motif bears resemblance to the RHIM and RHIM-like motifs controlling necroptosis in mammals and innate immunity in flies. We identified here, a third NLR signalosome in *Podospora anserina* comprising a PP-motif and organized as a two-gene cluster encoding a NLR and a HELL-domain cell-death execution protein termed HELLP. We show that the PP-motif region of HELLP forms a prion we term [π] and that [π] prions trigger the cell-death inducing activity of full length HELLP. We detect no prion cross-seeding between HET-S, HELLF and HELLP amyloid motifs. In addition, we find that akin to PP-motifs, RHIM motifs from human RIP1 and RIP3 kinases are able to form prions in *Podospora*, and that [π] and [Rhim] prions partially cross-seed. Our study shows that *Podospora anserina* displays three independent cell-death inducing amyloid signalosomes. Based on the described functional similarity between RHIM and PP, it appears likely that these amyloid motifs constitute evolutionary related cell-death signaling modules.

**Importance:** Amyloids are β-sheet-rich protein polymers that can be pathological or display a variety of biological roles. In filamentous fungi, specific immune receptors activate programmed cell-death execution proteins through a process of amyloid templating akin to prion propagation.

Among these fungal amyloid signaling sequences, the PP-motif stands out because it shows similarity to RHIM, an amyloid sequence controlling necroptotic cell-death in mammals. We characterized an amyloid signaling system comprising a PP-motif in the model species *Podospora anserina* thus bringing to three the number of independent amyloid signaling cell death pathways described in that species. We then show that human RHIM motifs not only propagate as prions in *P. anserina* but also partially cross-seed with fungal PP-prions. These results indicate that in addition to show sequence similarity, PP and RHIM-motif are at least partially functionally related, supporting a model of long-term evolutionary conservation of amyloid signaling mechanisms from fungi to mammals.

## Introduction

Prions are protein polymers that behave as infectious entities by propagating their amyloid structural state (1, 2). In addition to disease causing prions in mammals, prions have been identified in fungi as infectious, cytoplasmic, non-mendelian genetic elements (3). One such prion is [Het-s] of the filamentous fungus *Podospora anserina.* [Het-s] functions in a non-self recognition process known as heterokaryon incompatibility. The *het-s* gene exists as two incompatible allelic variants termed *het-s* and *het-S*. The HET-s protein when assembled into the self-propagating prion form confers incompatibility to the HET-S variant. Co-expression of the [Het-s] prion with HET-S leads to cell death. At the macroscopic level, the incompatibility reaction leads to the formation of an abnormal contact line between the strains termed “barrage”. The soluble non-prion form confers a phenotype termed [Het-s*] compatible with HET-S (4, 5). [Het-s*] strains acquire the prion state either spontaneously at a low frequency or systematically after fusion with a prion infected strain. HET-s and HET-S are highly homologous allelic variants that display two domains, a C-terminal Prion-Forming Domain (PFD) necessary and sufficient for prion propagation and a N-terminal α-helical globular domain named HeLo responsible for cell-death inducing activity (6, 7). The HET-s PFD is natively unfolded in the soluble state and upon prion formation assembles into to a specific cross-β amyloid structure (6, 8–10). The HET-s β-solenoid fold is composed of 2 rungs of β-strands per monomer, comprising two 21 amino acid long imperfect repeats (R1 and R2) connected by a flexible loop (9, 11). In [Het-s]/HET-S incompatibility cell death is triggered when the [Het-s] PFD templates conversion of the HET-S PFD region into the β-solenoid fold which in turn induces refolding of the HET-S HeLo domain that acquires pore-forming activity by exposing an N-terminal transmembrane helix targeting the cell membrane (7, 12). In this incompatibility system, HET-S acts as cell-death execution protein and [Het-s] triggers its activation.

Further work on the [Het-s] system revealed that this incompatibility system is evolutionary derived from a programmed cell death (PCD) pathway in which HET-S is activated by a specific NLR (Nod-like receptor) (13). NLR receptors are intracellular receptors that control immune defense and PCD pathways in animals, plants and fungi and function by ligand induced oligomerization (14, 15). The NWD2 NLR receptor is encoded by the gene immediately adjacent to *het-S* in the *P. anserina* genome and displays a central NACHT nucleotide binding and oligomerization domain and a C-terminal WD40 repeat domain. The N-terminal region of NWD2 displays a short region of homology to the R1 and R2 motifs of HET-s. This region designated R0 is able to adopt a *het-s*-like fold. Ligand induced oligomerization of NWD2 allows for spatial clustering of R0 motifs, cooperative nucleation of the β-solenoid fold which then templates conversion of the homologous domain in HET-S and activation of the HeLo domain (13, 16, 17). More generally, it is proposed that a fraction of the NLR receptors in fungi activate their cognate effector domains by a mechanism of prion amyloid templating by which the amyloid fold formed after oligomerization of the NLR receptor is transmitted through heterotypic interactions to the amyloid motif found in the C-terminal of the effector protein (18). NWD2/HET-S pair was used as a paradigm in bioinformatic screens to identify additional NLR/effector pairs functioning through amyloid templating in fungi (16, 19). These approaches resulted in particular in the identification of five *het-s*-related amyloid motifs families (HRAMs) and in the characterization in *Podospora anserina* of a distant homolog of HET-S termed HELLF and belonging the HRAM5 family (20). HELLF encodes a protein with a HeLo-like N-terminal domain and a C-terminal PFD with two repeated submotifs that share only 17% identity to the HET-s PFD. The HeLo-like domain shows distant homology to the HET-S HeLo domain but also to the 4HB pore-forming domain of the MLKL protein (21), the terminal effector protein of necroptotic death in mammals, which led to the proposition that this form of fungal PCD is related to mammalian necroptosis (22). The gene adjacent to *hellf* encodes a NLR with a NB-ARC domain and TPR repeats termed FNT1, which bears an N-terminal R0 region homologous to the HELLF R1 and R2 PFD repeats. HELLF behaves analogously to HET-S in every aspect that was analyzed. The C-terminal region of HELLF forms a prion termed [Φ] and [Φ] is incompatible with full length HELLF. Upon interaction with [Φ], HELLF relocates to cell membrane region and causes cell death. The N-terminal region of FNT1 forms fibrils and is able to induce [Φ] prion formation. The solid-state NMR structure of the HELLF PFD structure revealed that HET-s and HELLF prions have an almost identical backbone structure in spite of extensive sequence divergence. The FNT1/HELLF system thus constitutes a second cell death-inducing amyloid signaling pathway of *Podospora anserina*. Noteworthy is the fact that in spite of their almost identical backbone structure [Het-s] and [Φ] prions do not cross-seed indicating that NWD2/HET-S and FNT1/HELLF constitute two parallel signaling pathways.

In bioinformatics surveys, additional amyloid signaling motifs were identified in other fungi (14, 16). In particular, a prion motif termed PP was identified in the genome of *Chaetomium globosum* in a three-gene cluster and functionally studied using heterologous expression in *Podospora anserina*. In this specific case, the NLR termed PNT1 regulates activation of two effector proteins, a HeLo-like domain cell-death inducing protein termed HELLP and a putative lipase, SBP (renamed herein CgHELLP, CgPNT1 and CgSBP for clarity). The PP-motif is related in sequence to the RHIM amyloid motif that controls assembly of the RIP1 and RIP3 kinases in the necroptosis pathway in mammals (13, 22, 23). RHIM-like amyloid motifs also occurs in the PGRP-LC, PGRP-LE, and Imd proteins controlling anti-bacterial immunity in Drosophila (24). The structure of the mammalian RIP1-RIP3 core (forming an hetero-amyloid signaling complex) has recently been established (25). RHIM amyloids comprise two protein units per cross-β layer that run antiparallel and are interdigitated in a compact hydrophobic interface formed by the core G-(I,V)-Q-(I,V)-G motif. This central core motif is common to the PP-motif but in absence of structural characterization of PP-amyloids, it cannot at present be ascertained that this short sequence homology reflects structural similarity between RHIM and PP-amyloids.

Herein, we mine the genome of *P. anserina* and identify HELLP, a novel PP-motif protein that defines, in that species, a third cell-death inducing amyloid signaling pathway comprising the HELLP cell-death execution protein and a NLR (PNT1). We show that the C-terminal PP-motif of HELLP (HELLP(214-271)) forms a prion termed [π] in *Podospora anserina* and assembles into fibrils *in vitro.* We find that akin to HET-S and HELLF, cell-death inducing activity of HELLP is activated by [π] prions formed by HELLP(214-271) or by the N-terminal region of PNT1, PNT1(1-31). We analyze in *Podospora anserina* the functional interactions of the HET-s, HELLF and HELLP PFD prions and find that they form three independent signaling systems. We further analyze prion formation by human RHIM motifs from RIP1 and RIP3 kinases and find that [Rhim] and [π] prions partially cross-seed underlying the functional similarity between these mammalian and fungal amyloid motifs.

## Results

### A PP-motif gene cluster in *Podospora anserina*

During the course of a survey of NLR encoding genes in fungi, we identified in *Podospora anserina*, a gene encoding a NLR with a NB-ARC domain and TPR repeats (*Pa_5_8060*, PNT1), (14). We found after manual re-annotation, that the adjacent gene (*Pa_5_8070*) encodes a HeLo-like domain protein, we term HELLP (Fig. 1A). HELLP shows 56 % similarity to the CgHELLP cell-death execution protein of *Chaetomium globosum* (Fig. 1B and Fig. S1) (22). As previously reported for CgHELLP and HELLF, the HELL domain of HELLP shows homology to the 4HB membrane-targeting domain of the mammalian MLKL necroptosis execution protein (Fig. S1). The N-terminal region of PNT1 and the C-terminal region of HELLP share a region of homology encompassing a predicted PP-amyloid motif (Fig. 1B) (22). This region is also homologous to the RHIM motif found in the RIP1 and RIP3 kinases in humans (23, 25). Based on the resemblance with the PP gene-cluster of *C. globosum*, we reasoned that after *nwd2/het-s* and *fnt1/hellf*, the *pnt1/hellp* gene pair might encode the components of a third amyloid NLR signalosome in Podospora (Fig. 1A). We thus engaged into the functional characterization of this gene pair.

**Fig 1.**
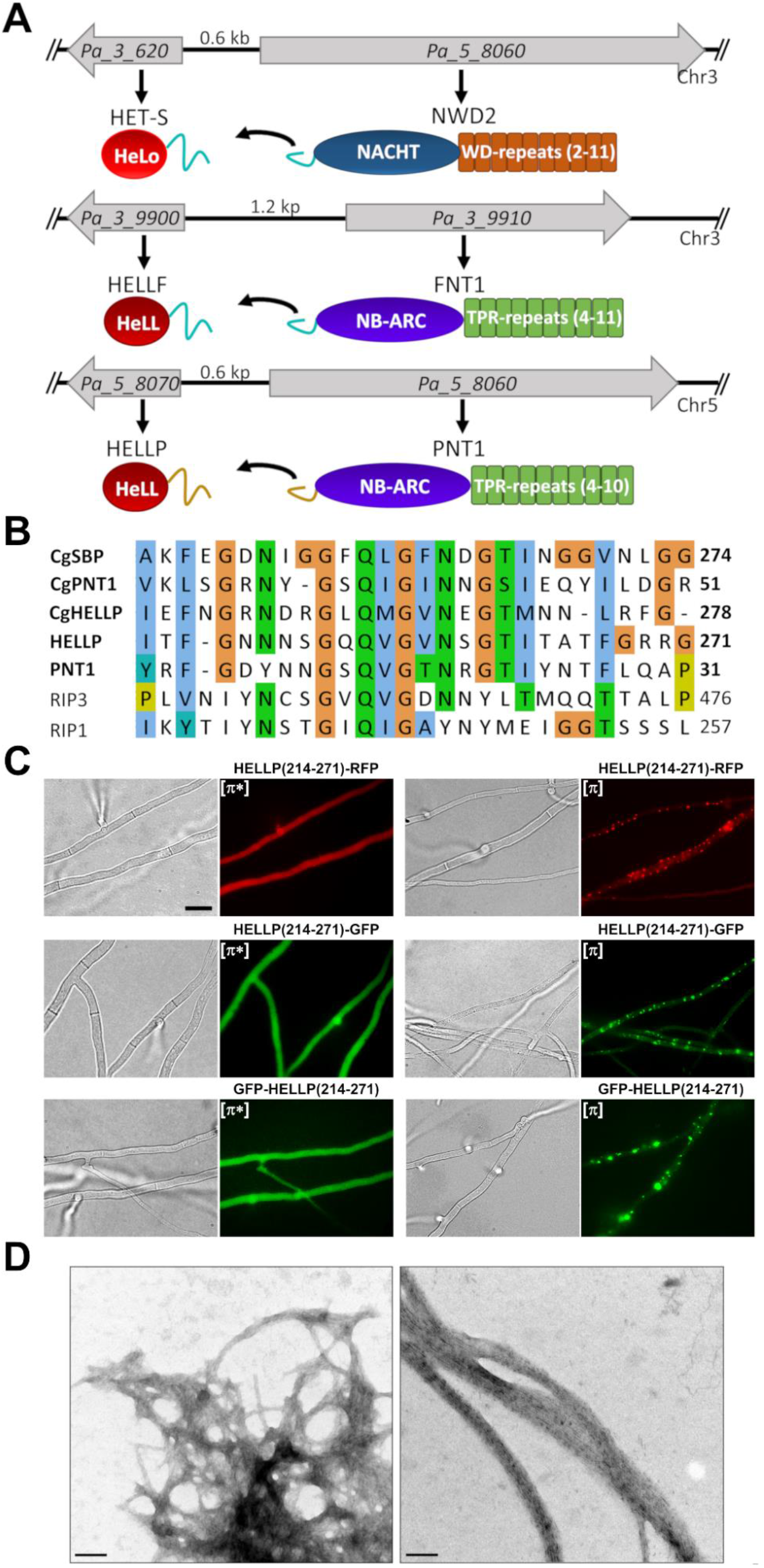
Characterization of the PP motif of *P. anserina.* (A) Genome organization and domain architecture of the *het-s/nwd2*, *hellf/fnt1* and *hellp/pnt1* gene clusters of *P. anserina*, chromosome, gene orientation and intergenic distances are given. These loci are composed of two adjacent genes encoding an effector protein and a NLR receptor. The effector proteins are composed of a PFD and a HeLo or HELL domain. The NLRs are composed of a PFD, a NB-ARC or NACHT nucleotide binding domain and of WD or TPR repeats (repeat numbers are variable in wild-isolates as indicated). PFDs of the HRAM family given in cyan and those of the PP family in light-brown line. (B) Alignment of the PP motifs of HELLP, PNT1, CgHELLP, CgSBP, and CgPNT1 along with the RHIM motif of human RIP1 and RIP3. (C) PP-motif region of HELLP behaves as a prion forming domain *in vivo* as shown on micrographs of *P. anserina* strains expressing different molecular fusions (HELLP(214-271-RFP/GFP, GFP-HELLP(214-271)) as indicated above each micrograph, (scale bar is 5 μm). Transformants present initially a diffuse fluorescence in the non-prion state designated [π*] (left) and systematically acquire dot-like fluorescent aggregates after contact with a strain already expressing the infectious prion state designated [π] (right). (D) PP-motif forms fibrils *in vitro* as shown on electron micrograph of negatively stained HELLP(214-271) fibrils, (scale bar is 100 nm).

### The C-terminal region of HELLP is a prion-forming domain

In order to analyze whether the C-terminal region of HELLP displays the anticipated prion forming ability, we generated different constructs with the C-terminal domain of HELLP (214-271) fused either to GFP or RFP (in a C-terminal position) or to GFP in an N-terminal position. These constructs were expressed in a *Δ*hellp strain of *P. anserina*. For the different constructs, a population of 25 to 47 distinct transformants was analyzed by fluorescence microscopy. We observed either dot-like or diffuse fluorescence (Fig. 1C). The fraction of transformants exhibiting dot-like fluorescence was found to increase with growth duration (Table 1). The highest rate of foci formation was observed for the fusion protein bearing the GFP in N-terminal position. Within 19 days, all tested GFP-HELLP(214-271) transformants, displayed fluorescent foci. For all three fusion constructs, strains with diffuse fluorescence were systematically converted to the foci phenotype after cytoplasmic contact with strains expressing foci (Table 1), indicating that the foci state is transmitted by cytoplasmic contact and infectious. By analogy with the [Het-s] system, we term the phenotypic state with diffuse fluorescence [π*] and the foci state [π] (Fig. 1C). We conclude from these experiments that the PP-motif region of HELLP(214-271) allows for prion formation and propagation. To determine whether as previously described for [Het-s] (26), the [π] state could be reverted to [π*] in meiotic progeny, we analyzed 40 progeny of a *Δhellp Δhet-s Δhellf GFP-hellp(214-271)* [π] × *Δhellp Δhet-s Δhellf het-s-RFP* cross. Among the 20 progeny containing the *GFP-hellp(214-271)* transgene, three displayed a [π*] phenotype, indicating that akin to [Het-s], the [π] prion can be cured in sexual crosses.

**Table 1.**
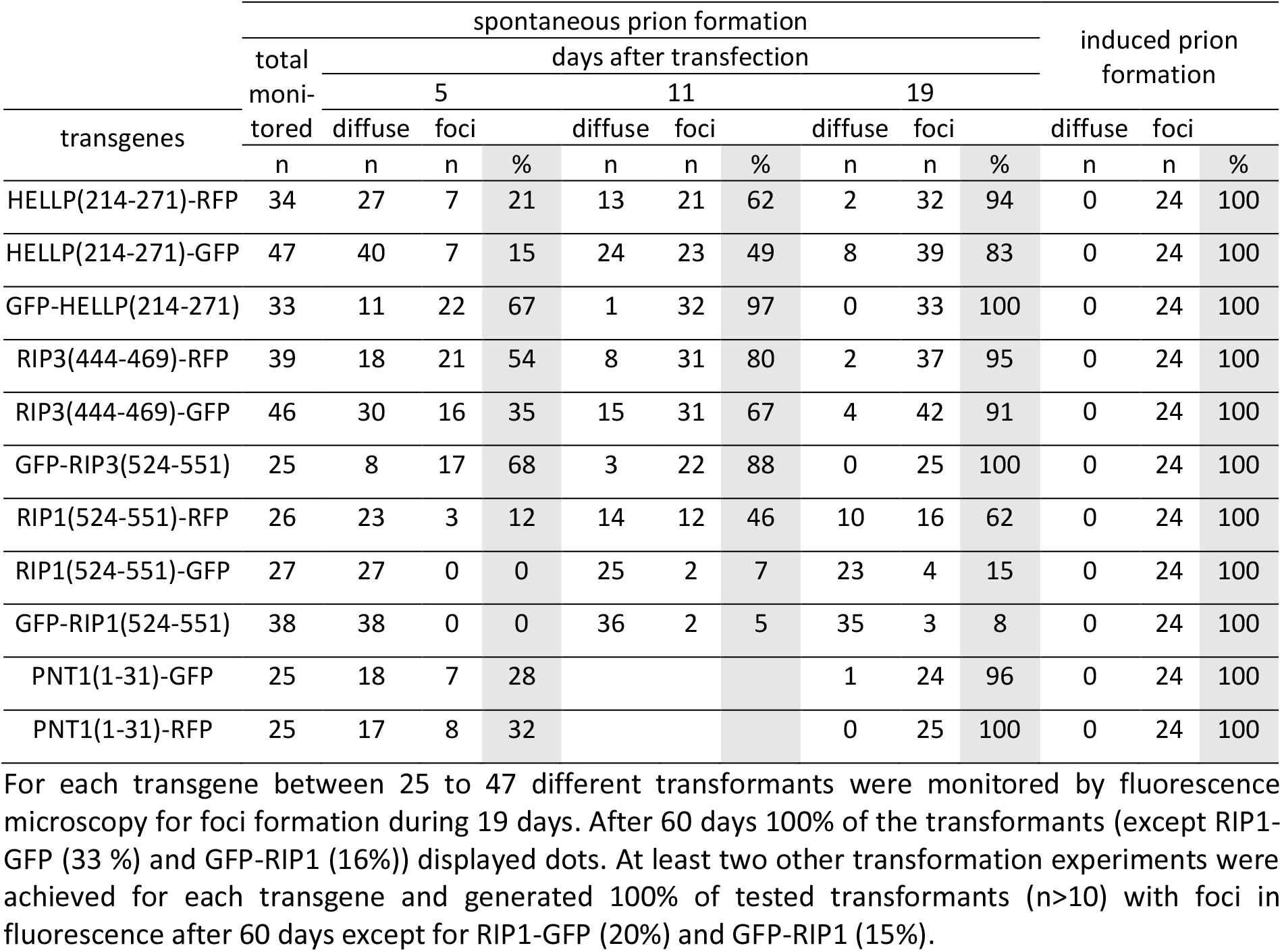
Rate of spontaneous and induced prion formation and propagation.

A larger construct, with HELLP(171-271) fused to GFP either in N- or C-terminus also led to foci formation (Fig. S2A) but in contrast to analogous HET-S and CgHELLP constructs (HET-S(157-289)-GFP and CgHELLP(170-278)-GFP), HELLP(171-271) fusions did not form elongated aggregates (22, 27).

The HELLP(214-271) PFD region was also expressed in *E. coli* with a C-terminal histidine tag and purified under denaturing conditions. Upon removal of the denaturant by dialysis, the protein spontaneously formed fibrils that often associated laterally as bundles (Fig. 1D).

### [π] prions triggers incompatibility upon interaction with full-length HELLP

It was shown that HET-S, HELLF and CgHELLP cell-death inducing activity is triggered by interaction with the prion form of their respective PFDs (7, 12, 20, 22). As a result, confrontation of strains expressing PFDs in the prion state with strains expressing the corresponding full-length protein leads to an incompatibility reaction which results in death of the fusion cells and, at the macroscopic level, in formation of an abnormal contact line termed barrage (20, 22, 27, 28). To determine whether HELLP could also determine incompatibility, strains expressing GFP-HELLP(214-271) were confronted to strains expressing full-length HELLP, either from the wild-type resident copy or from HELLP-GFP or HELLP-RFP transgene copies (in a *Δ*hellp background) (Fig. 2A). A barrage reaction was observed systematically and specifically in confrontations with strains expressing the [π] phenotype (as determined by presence of foci). [π*] strains did not produce a barrage reaction. 24 transformants were analyzed in parallel for barrage formation and presence of foci and the two phenotypes were strictly correlated. The barrage reaction was stronger with strains expressing HELLP-GFP or -RFP fusions from the strong *gpd* promoter than with strains expressing HELLP from the resident copy as shown in Fig. 2A. We verified that at the microscopic level, barrage formation was indeed associated with cell death using the vital dye methylene blue (Fig. S4).

**Fig 2.**
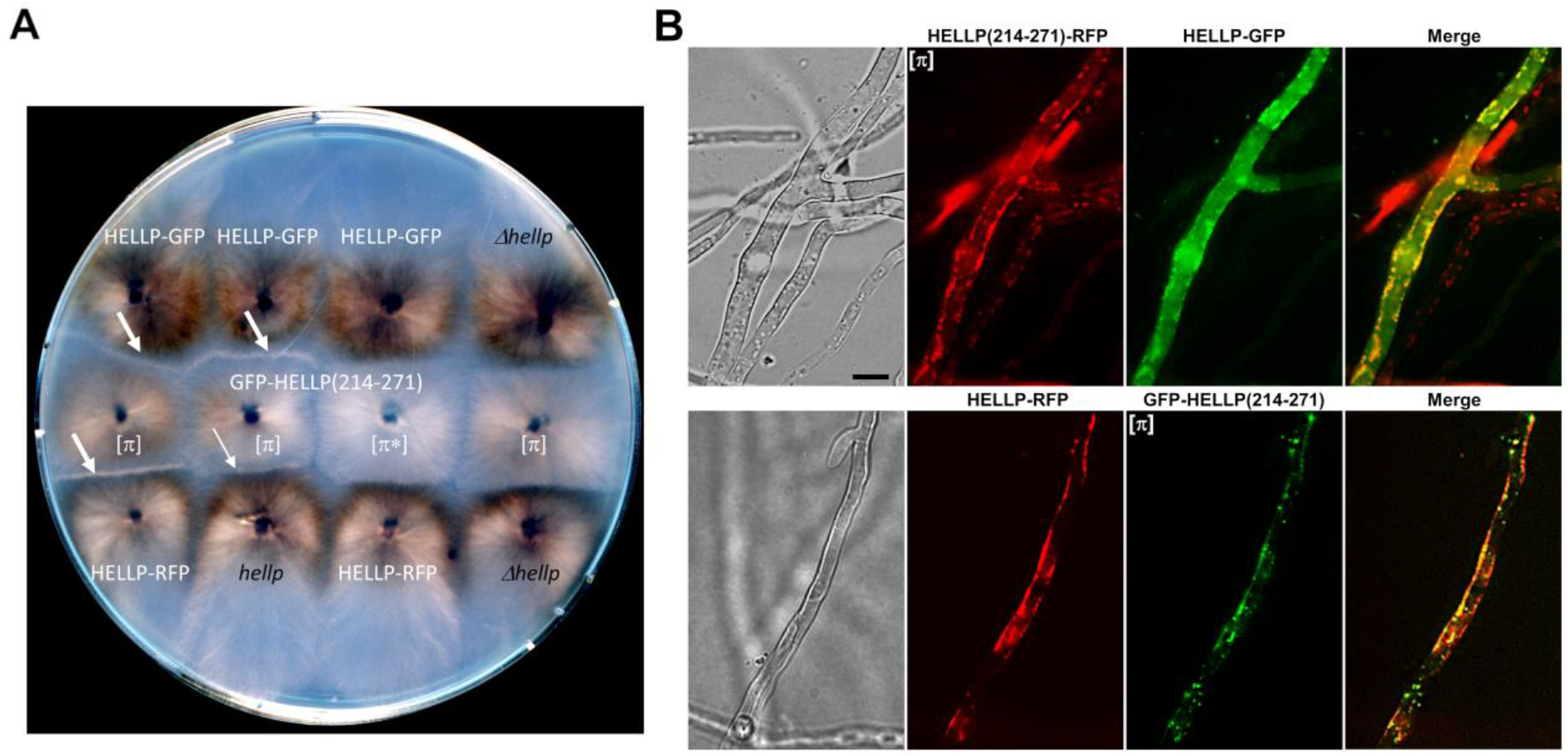
Activation of HELLP cell-death inducing activity by [π] prions. (A) Observation of incompatibility reaction (barrage) between a strain containing [π] prions and expressing HELLP. Confrontation on solid medium of transformants expressing GFP-HELLP(214-271) in soluble [π*] or aggregated [π] states with strains expressing full-length HELLP from a transgene (labelled in white) or from the wild type *hellp* gene or with the *Δhellp* strain (both labelled in black). Barrages indicative of an incompatibility reaction, are marked by white arrows. The thinner arrow indicates an attenuated barrage reaction. (B) Micrographs of fusion cells co-expressing full-length HELLP (HELLP-GFP (upper panel) or HELLP-RFP (lower panel)), and HELLP(214-271)-RFP (upper panel) or GFP-HELLP(214-271) (lower panel) in the [π] aggregated state. Note that HELLP relocalizes to the cell membrane region in presence of [π], (scale bar is 5μm).

As expected the barrage formation phenotype was transmitted by cytoplasmic contact. Upon contact with barrage forming strains, a compatible strain invariably became capable of forming a barrage with a strains expressing full length HELLP (Tables S1 and also S2 to S6, Fig. S2). Based on these experiments the definition of the [π] phenotype can be expanded to represent both the foci state of the PP-motif GFP and RFP fusions and the ability to produce a barrage reaction to full-length HELLP. The [π*]/[π] phenotypes of HELLP(214-271) are in that sense analogous to [Het-s*]/[Het-s] phenotypes of HET-s (4) and [Φ*]/[Φ] phenotypes of HELLF(209-277) (20).

It was shown that prion conversion of the HET-S PFD leads to the refolding of the HET-S HeLo domain that acquires pore-forming activity by exposing its N-terminal transmembrane helix, relocating it to the cell membrane and inducing cell death (7, 12). The same relocalization process was previously observed for HELLF and CgHELLP in the incompatibility reaction leading to cell death (20, 22). We thus determined whether HELLP relocates in the cell membrane region upon interaction with [π] prions. We examined the contact zone between incompatible [π] and HELLP-GFP or -RFP expressing strains (Fig. 2B). While full length HELLP-GFP or -RFP alone yield a diffuse fluorescence signal that remains stable over time (Fig. S3), in fusion cells co-expressing these proteins and the prion form of HELLP(214-271)-RFP or GFP-HELLP(214-271), we observed a relocalization of HELLP in the vicinity of the cell membrane region. We conclude that in these experiments, HELLP behaves as previously described for HET-S, HELLF and CgHELLP and locates in the cell membrane region in fusion cells. Further experiments are required to determine whether HELLP directly interacts with the membrane and if so how this interaction causes cell death.

### The PNT1(1-31) region forms a prion and induces HELLP activation

It has been shown that the N-terminal region of NWD2 adopts an amyloid fold, displays prion infectivity and is able to activate HET-S pore-forming activity (13). Similarly, N-terminal regions of the FNT1 and CgPNT1 NLRs behave as PFDs and trigger cell-death inducing activity of their cognate effectors proteins, HELLF and CgHELLP respectively (22). To determine whether the N-terminal region of PNT1 displays similar properties, we expressed PNT1(1-31) fused to GFP or RFP in *Δhellp* strains. Two phenotypic states were observed, a diffuse cytoplasmic fluorescence state, and a foci state analogous to the [π*] and [π] phenotypes observed previously with HELLP(214-271) (Fig. 3A). Spontaneous conversion to the foci form occurred upon prolonged subculture (Table 1) and cytoplasmic contact with foci-containing strain induced systematic conversion to the foci state (Table 1). PNT1(1-31)-GFP strains with fluorescent foci produced a barrage reaction to strains expressing full length HELLP (not shown). PNT1(1-31)-GFP [π] strains systematically convert [π*] HELLP(214-271)-RFP strains to the [π] state. Conversely, [π] HELLP(214-271)-RFP systematically convert [π*] PNT1(1-31)-GFP to the [π] state (Table S2).

**Fig 3.**
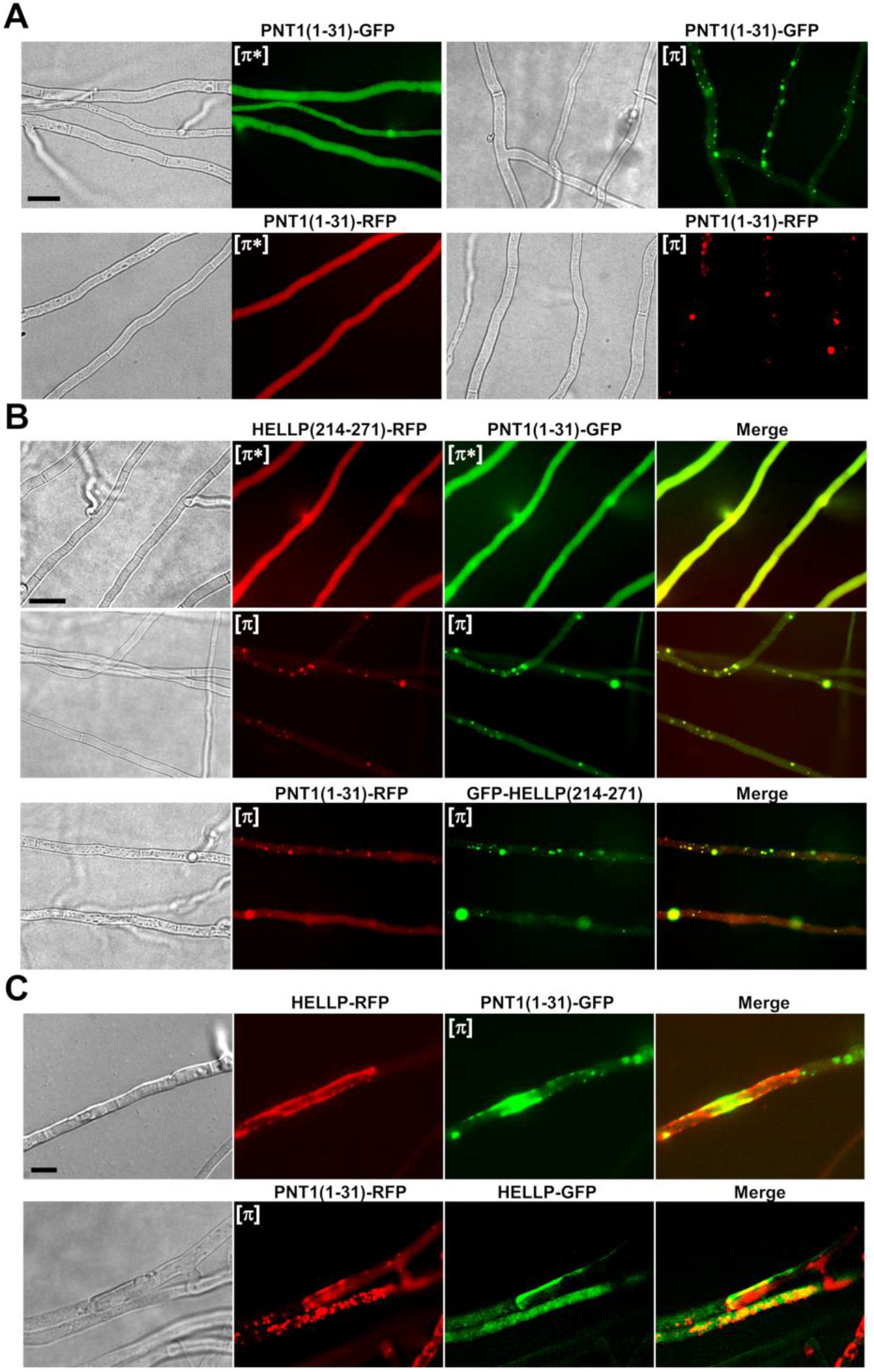
PNT1(1-31) forms [π] prions, co-localizes with HELLP(214-271) and induces toxicity of full-length HELLP. (A) Micrographs of PNT1(1-31) fusions with GFP or RFP either in the diffuse [π*] (left) or foci [π] state (right). (B) Micrographs showing co-localization of PNT1(1-31) and HELLP(214-271) in diffuse (top picture) or foci states. (C) In fusion cells co-expressing full-length HELLP (fused with RFP (upper panel) or GFP (lower panel)), and foci forms of PaPNT1(1-31) (fused with GFP (lower panel), or RFP, lower panel), HELLP relocalized to the plasma membrane region, (scale bar is 2 μm).

PNT1(1-31) and HELLP(214-271) were then co-expressed in the same strain (Fig. 3B). Fluorescence was either diffuse for both HELLP(214-271)-RFP and PNT1(1-31)-GFP or dot-like concomitantly for the two proteins. Cells in which one of the proteins formed foci while the other remained diffuse were not observed (Fig. 3B, Table S7). HELLP(214-271)-RFP and PNT1(1-31)-GFP dots generally co-localized (Fig. 3B, Table S8). The same was true during co-expression of PNT1(1-31)-RFP and GFP-HELLP(214-271). An initial diffuse state could however not be observed in this setting presumably as the result of the high spontaneous conversion rate of the N-terminal GFP-HELLP(214-271) fusion protein (Table 1).

Finally, we observed fusion cells between strains expressing PNT1(1-31) in the [π] state and strains expressing full-length HELLP (Fig. 3C, Fig. S4). We observed the relocalization of HELLP to the plasma membrane region and cell death of the fusion cells. Thus, consistent with the proposed role of HELLP as cell death execution protein activated by the PNT1 NLR, we find that the PP-motif region of PNT1 forms a prion and is able to activate HELLP.

### HET-S, HELLF and HELLP are components of three independent cell-death inducing pathways

We recently reported the lack of cross-seeding between [Het-s] and [Φ] prions indicating the existence of two independent parallel amyloid signaling pathways in *P. anserina* (20). Here, we identify HELLP as an effector in a third amyloid signaling pathway in *P. anserina* and therefore wanted to analyze potential cross-interaction between these pathways. We analyzed [π*] conversion by [Het-s] and [Φ] prions (Table S3). Strains expressing HELLP(214-271)-GFP or -RFP and displaying the [π*] phenotype were confronted to strains expressing [π], [Φ] or [Het-s] prions and then tested for their ability to form a barrage with tester strains expressing full-length HELLP. Barrage were observed only (and systematically) when the prion donor strain expressed [π] prions indicating that [π*] conversion does not occur in presence of [Het-s] or [Φ] prions.

We also analyzed the cross-activation of the cell-death effectors of the three systems by the different prions (Table S4). We used incompatibility assays and confronted [π], [Φ] or [Het-s] prion expressing strains with strains expressing either HELLP, HELLF or HET-S full-length proteins. We observed barrage formation solely when [π] strains were confronted with HELLP expressing strains, [Φ] with HELLF and [Het-s] with HET-S, that is, uniquely when the prion and the effector bear the same PFD. We observe no cross-interaction between these three amyloid signaling pathways.

To further analyze HELLP/HELLF and HELLP/HET-s interactions, we co-expressed HELLP(214-271) with either HET-s or HELLF(209-271) in a *ΔhellpΔhet-sΔhellf* strain. We observed the diffuse and foci states co-existing in all possible combinations (Fig. 4, Fig S5, and Table S7). Namely, strains expressing HELLP(214-271) and HET-s could display four different phenotypes: [Het-s]/[π] and [Het-s*]/[π*] but also [Het-s]/[π*] and [Het-s*]/[π]. The same was true for HELLP(214-271) and HELLF(209-277) co-expression (Fig. 4, S5 and Table S7). These results are consistent with the conversion experiments and show that [Φ] or [Het-s] prions do not efficiently convert [π*] and conversely that [π] prions do not efficiently convert [Φ*] or [Het-s*]. We also noted an absence of co-localization between HELLP(214-271) and HET-s or HELLP(214-271) and HELLF(209-277) prion forms (Fig. 4, Table S8).

**Fig 4.**
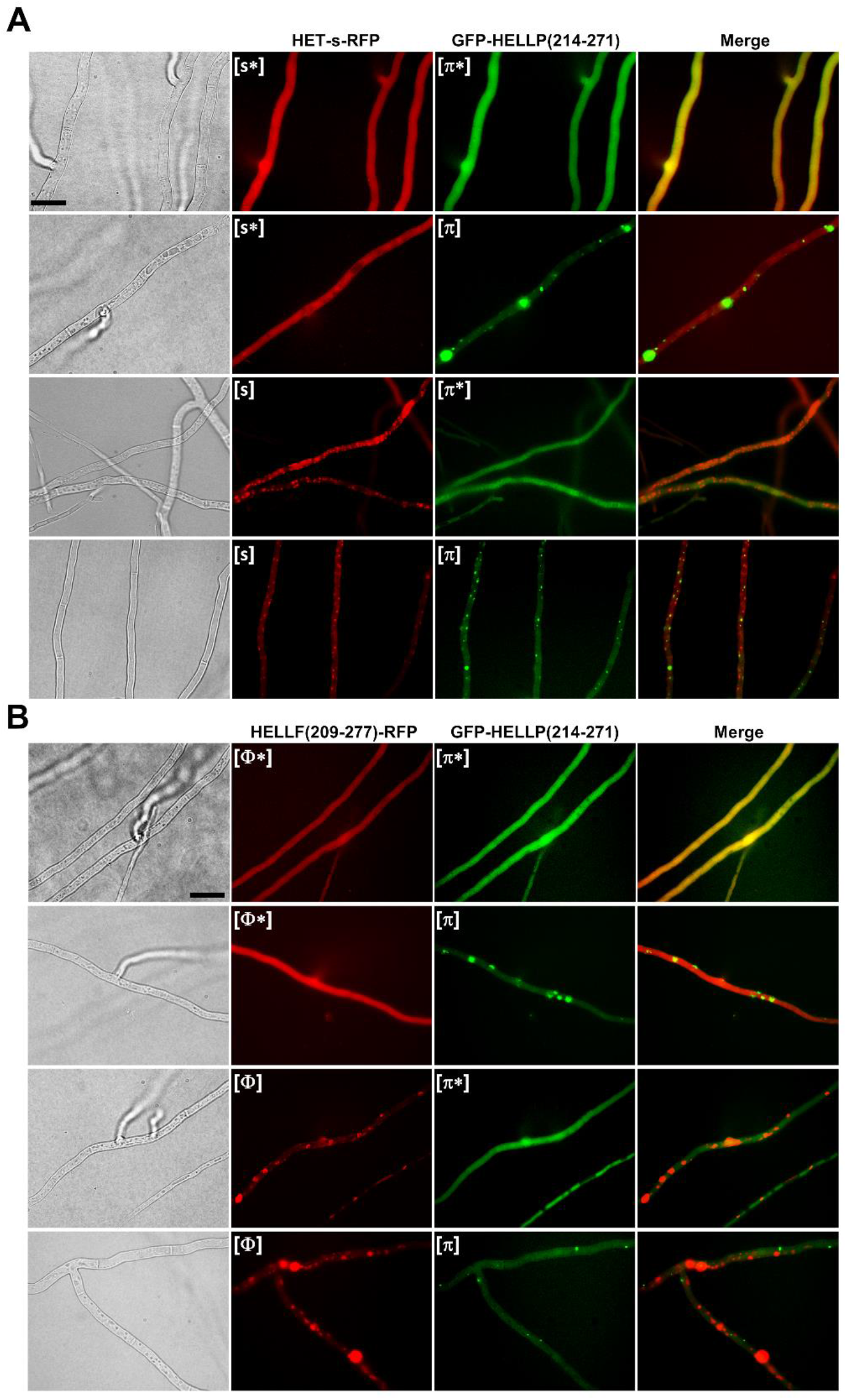
Lack of co-localization *in vivo* between [π] and [Het-s] or [Φ] prions. (A) Micrographs of strains co-expressing HET-s-RFP ([Het-s] or [Het-s*] states, noted here [s*] and [s]) and GFP-HELLP(214-271) ([π*] or [π] state). Note an absence of co-localization of the prion forms and the independent occurrence of [π] and [Het-s], (scale bar is 5 μm). (B) Micrographs of strains co-expressing HELLF(209-277)-RFP ([Φ] or [Φ*] state) and GFP-HELLP(214-271) ([π*] or [π] state). Note an absence of co-localization of the prion forms and the independent occurrence of [π] and [Φ] (scale bar is 5 μm), (scale bar is 5 μm).

### *P. anserina* and *C. globosum* [π] prions cross-seed and co-localize

It was shown previously that HET-S PFDs homologs from *P. anserina* and *Fusarium graminearum* which share about 41% identity do cross-seed (29). We wondered whether the 36% similarity between the PP-regions of HELLP and CgHELLP would also allow for cross-seeding (as suggested by the fact that pairwise similarities between the different PP-motifs within the *C. globosum* PP-gene cluster or within the *P. anserina* PP-gene pair are in the range of 32-48%, Fig. 1B).

Upon confrontation, [π] GFP-HELLP(214-271) donor strains invariably converted GFP-CgHELLP(215-278) [π*] recipient strains and conversely [π] CgHELLP(215-278) strains converted GFP-HELLP(214-271) [π*] recipients (Table S5). We analyze the cross-induction of cell death activity of the full-length CgHELLP and HELLP proteins in barrage tests (Fig. 5B and Tables S5, S6). GFP-CgHELLP(215-278) [π] strains produced a barrage reaction to strains expressing HELLP and conversely GFP-HELLP(214-271) [π] produced a barrage reaction to strains expressing CgHELLP. However, this was only true when HELLP was highly expressed as a transgene. No barrage was observed in confrontation to wild type. This result suggest that heterologous activation of HELLP by GFP-CgHELLP(215-278) is less efficient than homotypic activation by GFP-HELLP(214-271).

**Fig 5.**
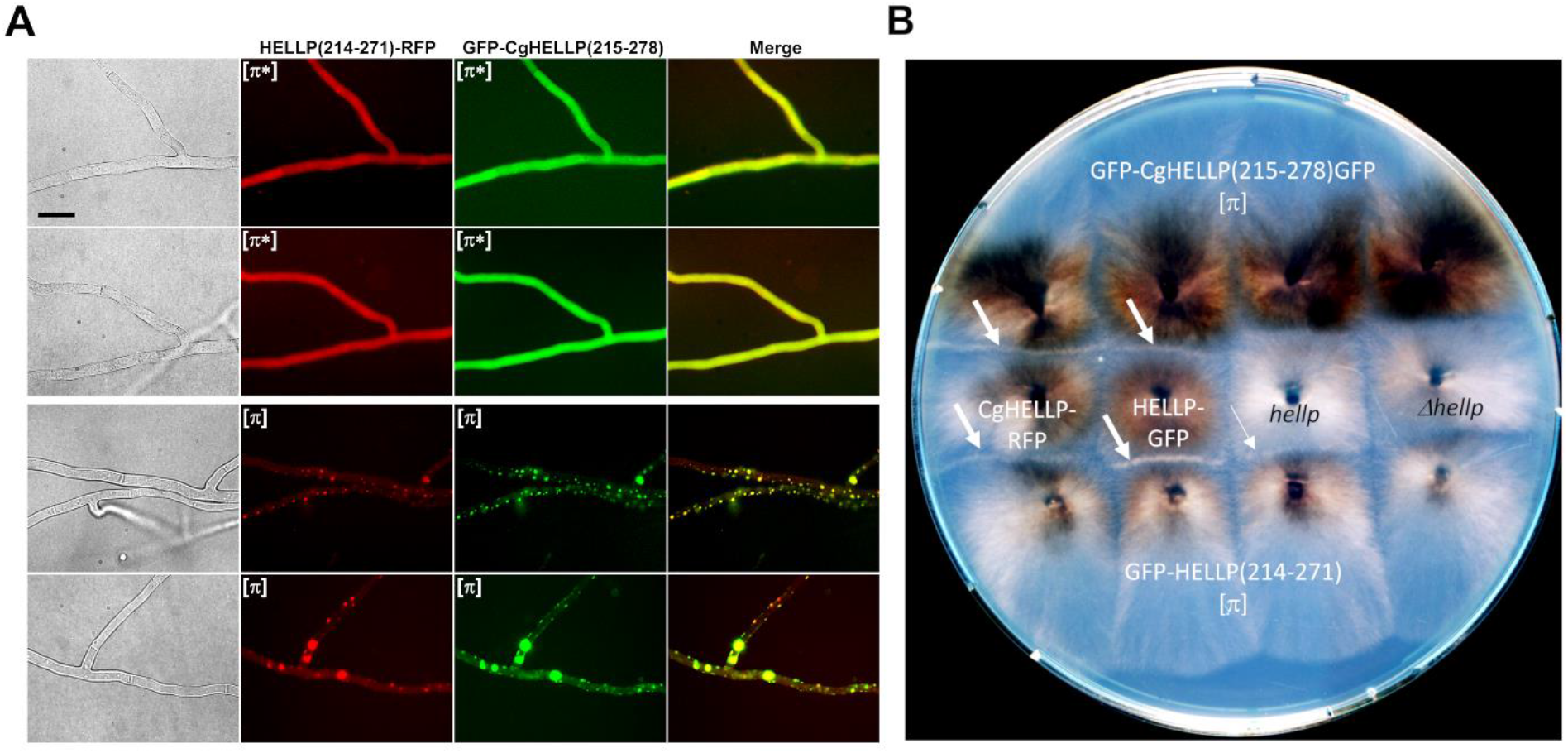
Co-localization of HELLP and CgHELLP PFDs and cross induction of cell-death activity. (A) Micrographs of strains co-expressing HELLP(214-271)-RFP and GFP-CgHELLP(215-278) in diffuse [π*] state (upper panels) or foci [π] state (lower panels). Note the co-localization of the two prion forms, (scale bar is 5 μm). (B) Confrontation on solid medium of strains expressing GFP-HELLP(214-271) and GFP-CgHELLP(215-278) [π] prions with strains expressing full-length HELLP from a transgene (marked in white) or from the wild type *hellp* gene or with the *Δhellp* strain (both marked in black). Barrages indicative of an incompatibility reaction, are marked by a white arrow. The thinner arrow indicates an attenuated barrage reaction.

To examine if the two PFD regions could co-localize *in vivo*, strains co-expressing HELLP(214-271)-RFP and GFP-CgHELLP(215-278) were obtained and analyzed by fluorescence microscopy (Fig. 5A, Tables S7 and 8). We initially observed diffuse fluorescence corresponding to the [π*] state and after a few days, foci appeared concomitantly for the two proteins. There was no co-existence of diffuse and foci forms suggesting cross-conversion of HELLP and CgHELLP (Table S7). Moreover, we observed a significant co-localization indicating co-aggregation of the two prions (Fig. 5A).

We conclude that, as previously reported for *F. graminearum* and *P. anserina* [Het-s], the “species-barrier” for prion seeding can be crossed for these PP-motifs. This result is consistent with the fact that the level of within-cluster similarity of the PP-motifs is comparable to the similarity between CgHELLP and HELLP PP-motifs. Nevertheless, the absence of barrage reaction between strains expressing [π]^Cg^ prions and HELLP at wild-type level suggests that homotypic or within-cluster interactions are more efficient to activate the cell-death activity of the full-length protein.

### The RHIM motifs of human RIP1 and RIP3 kinases form prions in *P. anserina*

The PP-motif as found in HELLP shows sequence homology to the mammalian RHIM amyloid motif (Fig. 1B). We thus envisioned that akin to the PP-motif, mammalian RHIMs might behave as PFDs when expressed in Podospora. We expressed the RIP1(524-551) and RIP3(444-469) regions fused to GFP or RFP in *ΔhellpΔhet-sΔhellf* strains (Fig. 6). Initially fluorescence was diffuse and cytoplasmic and foci appeared spontaneously after a few days of growth. The rate of spontaneous transition to the aggregated state was monitored as previously for HELLP (Table 1). For RIP3(444-469), the rates were comparable to the PP-motif with the highest rate observed with GFP in N-terminal position. For RIP1 RHIM, spontaneous rate of foci formation was lower especially for the two GFP fusions. The induced conversion through cytoplasmic infection (with the corresponding foci form) was equally effective for the RIP1 and RIP3 RHIM-motifs (Table 1). We designated the diffuse and infectious foci states [Rhim*] and [Rhim] respectively.

**Fig 6.**
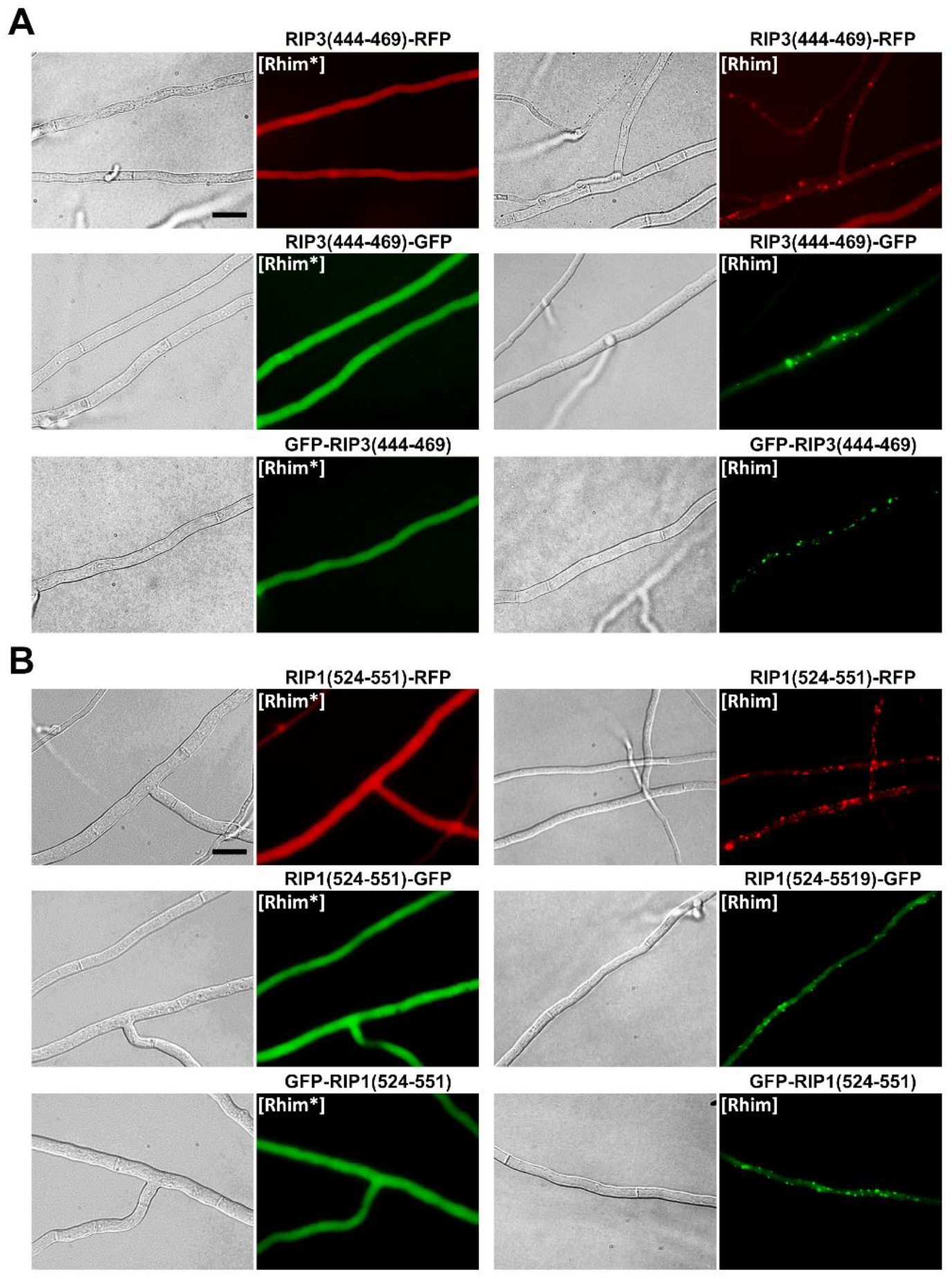
*In vivo* prion propagation of the human RIP3 and RIP1 RHIM regions in *Podospora anserina*. (A) Micrographs of *P. anserina* strains expressing the RIP3(444-469)-RFP, RIP3(444-469)-GFP or GFP-RIP3(444-469) molecular fusions in the diffuse ([Rhim*]) and foci state ([Rhim]), (scale bar is 5 μm). (B) Micrographs of *P. anserina* strains expressing the RIP1(524-551)-RFP, RIP1(524-551)-GFP or GFP-RIP1(524-551)molecular fusions in the diffuse ([Rhim*]) and foci state ([Rhim]), (scale bar is 5 μm).

Strains expressing RIP1(524-551) foci induced formation of RIP3(444-469) foci and the converse was true (Fig. 7) showing cross-conversions between [Rhim]^RIP1^ and [Rhim]^RIP3^ prions. In these assays, heterotypic cross-induction rates between [Rhim]^RIP1^ and [Rhim]^RIP3^ were not significantly different from homotypic induction rates. To further analyze the interaction of [Rhim] prions propagated in *P. anserina*, we co-expressed RIP1(524-551) and RIP3(444-469) fused to GFP or RFP in the same strain (Fig. 8, Tables S7 and S8). Strains co-expressing RIP3(444-469) fused with GFP and RFP, or RIP1(524-551) fused with GFP and RFP were used as positive control (Fig. 8B) and showed co-localization with overlapping foci. In RIP1(524-551) and RIP3(444-469) co-expression, diffuse fluorescence was observed initially. Then, foci formed concomitantly for both proteins. We observed no situation in which one of the protein formed foci while the other remained diffuse as previously observed for heterotypic PP-motif interactions (Fig. 8, Table S8). RIP1(524-551) and RIP3(444-469) foci co-localized in the cell (Fig. 8, Table S8), often in the close vicinity of the septa where they appeared as aligned dots.

**Fig 7.**
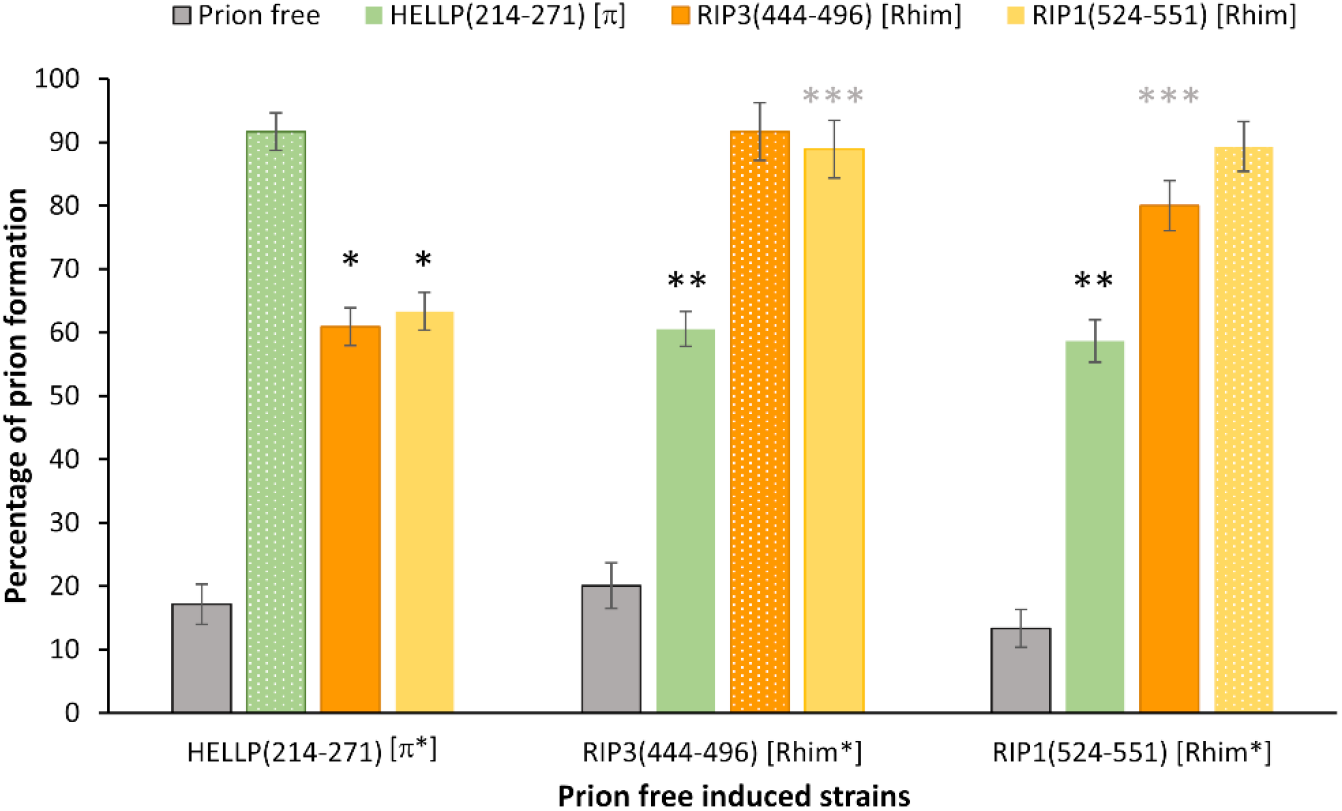
Cross conversion between HELLP(214-271) [π] and [Rhim] prions. Histograms representing the percentage of [π] and [Rhim] prion containing strains obtained after contact of recipient strains (initially displaying [π*] or [Rhim*] phenotype) with a *ΔhellpΔhet-sΔhellf* prion-free strain (control), a GFP-HELLP(214-271) [π] strain or GFP-RIP3(444-469) or GFP-RIP1(524-551) [Rhim] strains (as indicated on top). Phenotype after induction was determined by monitoring the acquisition of foci by fluorescence microscopy. Percentages of prion formation were expressed as the mean value ± standard deviation on 4 distinct transformants for each genotype (resulting in 56 to 108 independent infections per genotype). Homotypic inductions are showed with striped lines. P-values for cross conversion were determined using two tails Fisher’s test by comparison of the number of prion free and prion containing strains obtained after induction by the prion free control or by the heterotypic prion containing strain, (*: p-values <10^−8^, **: p-values <10^−12^).

**Fig 8.**
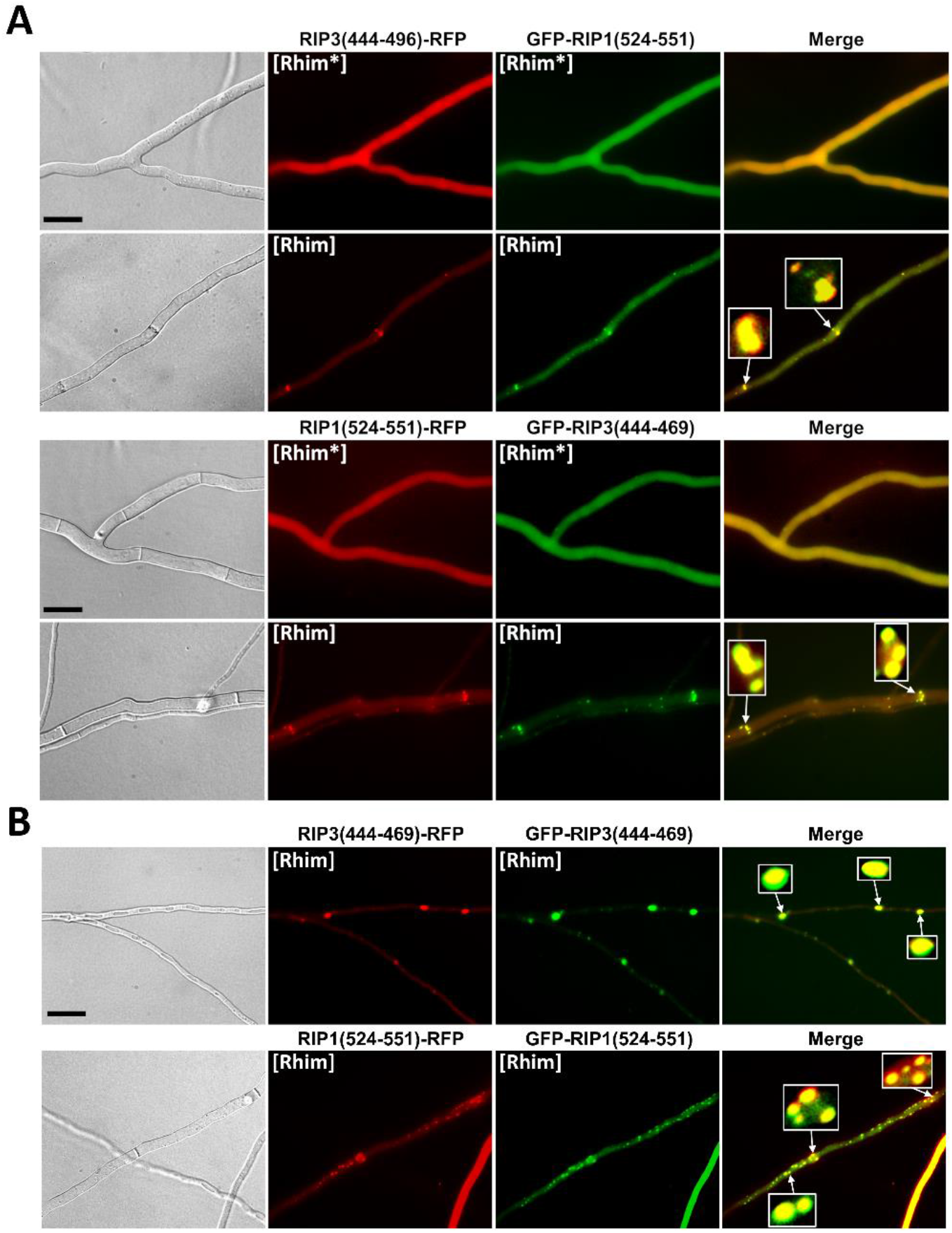
Co-localization of RIP1(524-551) and RIP3(444-469) foci in *Podospora anserina*. (A) Micrographs of *P. anserina* strains co-expressing the RIP3(444-469)-RFP and GFP-RIP1(524-551) or GFP-RIP3(444-469) and RIP1(524-551)-RFP molecular fusions, (scale bar is 5 μm). Note the strong co-localization between [Rhim] foci and dots alignment close to the septa (zoomed regions). (B) Micrographs of strains co-expressing GFP-RIP3(444-469) and RIP3(444-469)-RFP or GFP-RIP1(524-551) and RIP1(524-551)-RFP of [Rhim] phenotypes. Note the almost complete overlapping of foci of [Rhim] homopolymers, (scale bar is 5 μm).

We conclude that, the RHIM containing RIP3(444-469) and RIP1(524-551) regions of 26 and 28 amino acids in length behave as PFDs *in vivo* in *P. anserina* and lead to the expression of two alternate phenotypes we termed [Rhim*] and [Rhim]. In addition, the RIP1 and RIP3 [Rhim] prions efficiently cross-seed and co-localize consistent with the fact that in mammalian cells and *in vitro*, RIP1 and RIP3 interact through their RHIM regions to form amyloid heteropolymers (25).

### The [Rhim] and [π] prions partially cross-seed

Based on the similarity of PP and RHIM motifs and on the fact that RIP1 and RIP3 RHIM motifs propagate as prions *in vivo* in Podospora, it was conceivable that [π] and [Rhim] prions might cross seed to some extent. To examine possible interaction between [Rhim] and [π] prions, HELLP(214-271)-RFP [π*], GFP-RIP3(444-496) and GFP-RIP1(524-551) [Rhim*] strains were confronted with [π] and [Rhim] strains (Fig. 7). After four days, the recipient strains were sampled, subcultured, and analyzed for the presence of foci by fluorescence microscopy. We deliberately chose transformants expressing moderate levels of the fusion proteins to decrease the rates of spontaneous prion formation in these experiments. In these test conditions, spontaneous prion formation was in the range of 10 to 20% for all constructs. Homotypic interactions led to 80 to 90% prion conversion (as mentioned above, high conversion rates were observed for heterotypic conversion of GFP-RIP3(444-496) [Rhim*] by GFP-RIP1(524-551) [Rhim] and GFP-RIP1(524-551) [Rhim*] by GFP-RIP3(444-496) [Rhim]). [π] prions induced formation of both [Rhim] prions at a rate of about 60%. Conversely, both [Rhim] prions induced formation of [π] prions at a similar rate (Fig. 7). These results show cross conversion between [Rhim] and [π] prions. The conversion is however less efficient as in the case of homotypic or intra-kingdom [Rhim]^RIP1^/[Rhim]^RIP3^ or [π]^Pa^/[π]^Cg^ interactions.

We also tested the ability of the two [Rhim] prions to induce the conversion of [Het-s*] strains to the [Het-s] phenotype and observed cross-induction (18 transformants for each [Rhim] prion tested in triplicate). This result is in line with the previous results on the absence of cross-interaction between [π] and the other amyloid signaling pathways of *P. anserina* and are thus rather expected.

To explore interactions between [π] and [Rhim] further, HELLP and RIP3 (or RIP1) PFDs fused to GFP or RFP were co-expressed in the same strain and transformants were analyzed by fluorescence microscopy (Fig. 9, S6, Tables S7, S8). Consistent with the conversion experiments indicating partial cross-seeding, we could observe cells where one of the protein formed foci while the other remain in the diffused state as observed previously for non-cross seeding prions (Fig. 9 and S6, Table S7). In [π]/[Rhim] cells, the fraction of the foci that co-localized was higher (~68 %) than between non-cross seeding prions (such as HET-s and HELLP for instance, ~5 %) but lower than in homotypic or [Rhim]^RIP1^/[Rhim]^RIP3^ or [π]^Pa^/[π]^Cg^ prions (~82-96%). In addition, even in mixed foci the co-localization is imperfect with RFP and GFP fluorescence that only partially overlapped (Fig. 9). This phenomenon of patchy co-localization has already been observed during coexpression of the HET-s proteins of *F. graminearum* and *P. anserina* (29). The same [π]/[Rhim] interaction experiments were also carried out with CgHELLP(215-278) and similar results were obtained (Fig. S7, Tables S7 and S8). [π]^Cg^ and [Rhim] prions were found to partially cross-seed. And in this case also, there was a partial co-localization of [Rhim] and [π] prions, a situation that is distinct both from the total lack of interaction seen between [π] and [Het-s] and [Φ] prions and from the strong interactions seen between RIP1 and RIP3 [Rhim] prions on one hand and HELLP and CgHELLP [π] prions on the other hand.

**Fig 9.**
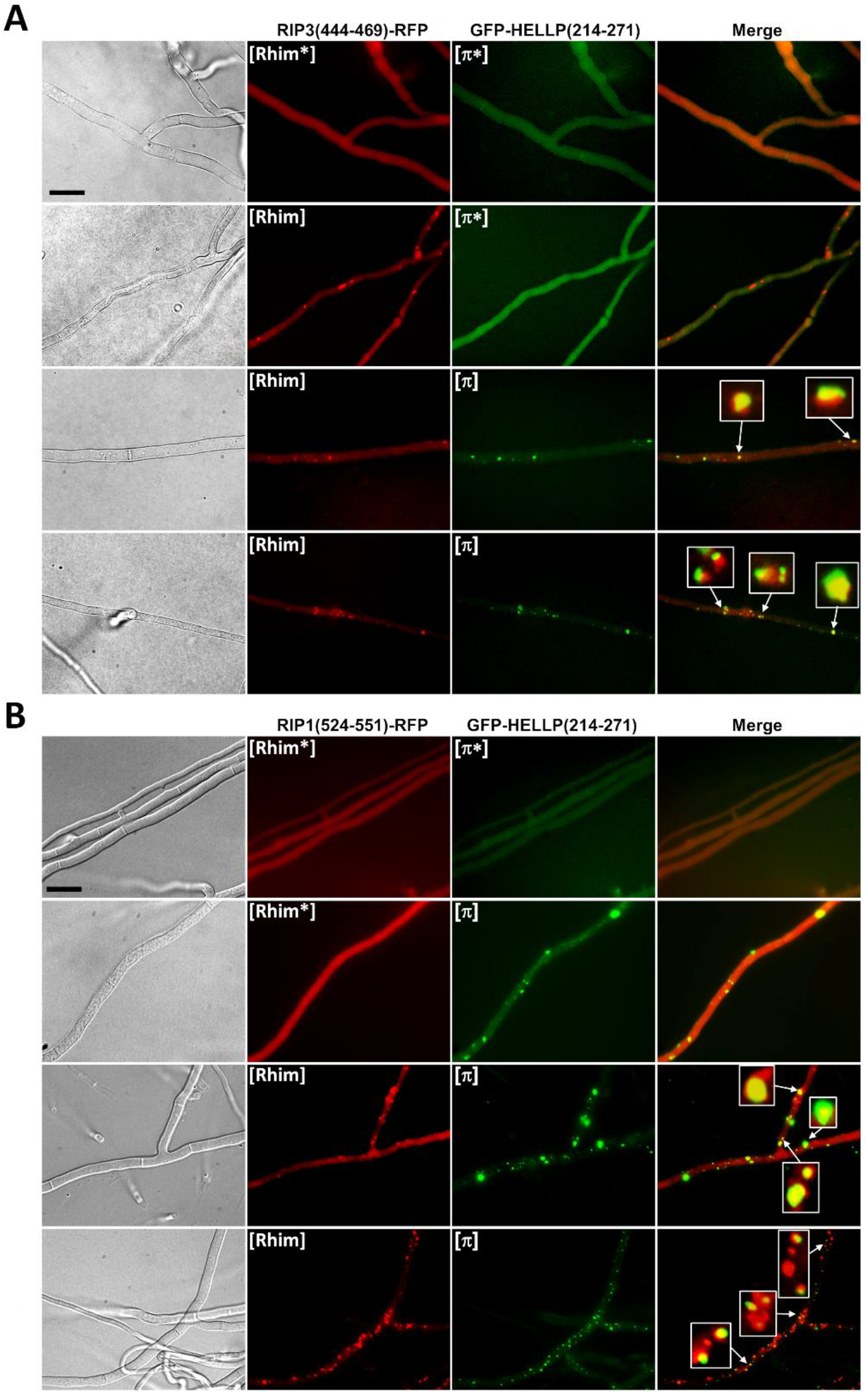
Partial co-localization of [π] and [Rhim] prions. (A) Micrographs of strains co-expressing RIP3(444-469)-RFP ([Rhim*] or [Rhim] states) and GFP-HELLP(214-271) ([π*] or [π] states). (B) Micrographs of strains co-expressing RIP1(524-551)-RFP ([Rhim*] or [Rhim] states) and GFP-HELLP(214-271) ([π*] or [π] states). In both cases, note the co-existence in the second lane of [Rhim] and [π*] (A) or [Rhim*] and [π] (B), these situations are only observable for a short period of time and upon prolonged subculture both prion forms occur together. Note also the partial co-localization of the two prions, some of the dots were zoomed to show incomplete overlapping in foci, (scale bar is 5 μm).

The above [Rhim]/[π] cross-seeding and co-localization experiments suggest that RHIM and PP-motifs can interact but that this interaction is less efficient than PP-homotypic interactions. We wondered whether [Rhim] prions could nonetheless induce at least partially, HELLP or CgHELLP toxicity in a setting favoring [Rhim]/HELLP interactions. We thus co-expressed in the same strain GFP-RIP3(444-469) or GFP-RIP1(524-551) and HELLP-RFP or CgHELLP to determine whether prion conversion of [Rhim] in this strain would lead to growth alterations (self-incompatibility). Positive controls corresponding to cross-interacting incompatible combinations ([π]/HELLP) or negative controls corresponding non-interacting combinations ([ϕ] or [Het-s]/HELLP or [Rhim]/HET-S) were used for comparison (Table 2). In each combination, growth of a population of 12 to 39 transformants was observed before or after infection with the respective prion. Before prion infection, growth was normal for the vast majority of the transformants. After prion infection, a large fraction of the transformants (about 80%) co-expressing RHIM constructs and full length HELLP or CgHELLP showed growth defects (Table 2 and Fig. 10). A similar proportion of growth defects was observed in known cross-interacting combinations while growth alterations where not significantly increased in non-interacting combinations. The growth alteration phenotypes were highly heterogeneous (Table2, Fig. 10) ranging from sublethal growth to minor growth alteration. This phenotypic heterogeneity is typical of such self-incompatible situations and was also observed for the control self-incompatible strains. This heterogeneity is thought to result from differences in transgene copy number and integration site in the different transformants and “escape” from self-incompatibility that is strongly selected for and often manifests as sector formation that recover close to normal growth and can occur by transgene mutation or deletion and in this particular case from prion curing. Strains with [Rhim] prions do not lead to a barrage reaction when confronted to strains expressing full length HELLP suggesting that the RHIM/HELLP interaction is not efficient enough to induce a massive cell-death reaction and barrage formation. We however observed cell death in fusion cells between strains expressing HELLP-RFP and GFP-RIP3(444-469) or GFP-RIP1(524-551) [Rhim] strains (Fig. S9). These results are consistent with the conversion experiments and indicate that [Rhim] prions are able to induce to some extent toxicity of HELLP (and CgHELLP) albeit at a lower efficiency than [π] prions.

**Table 2.**
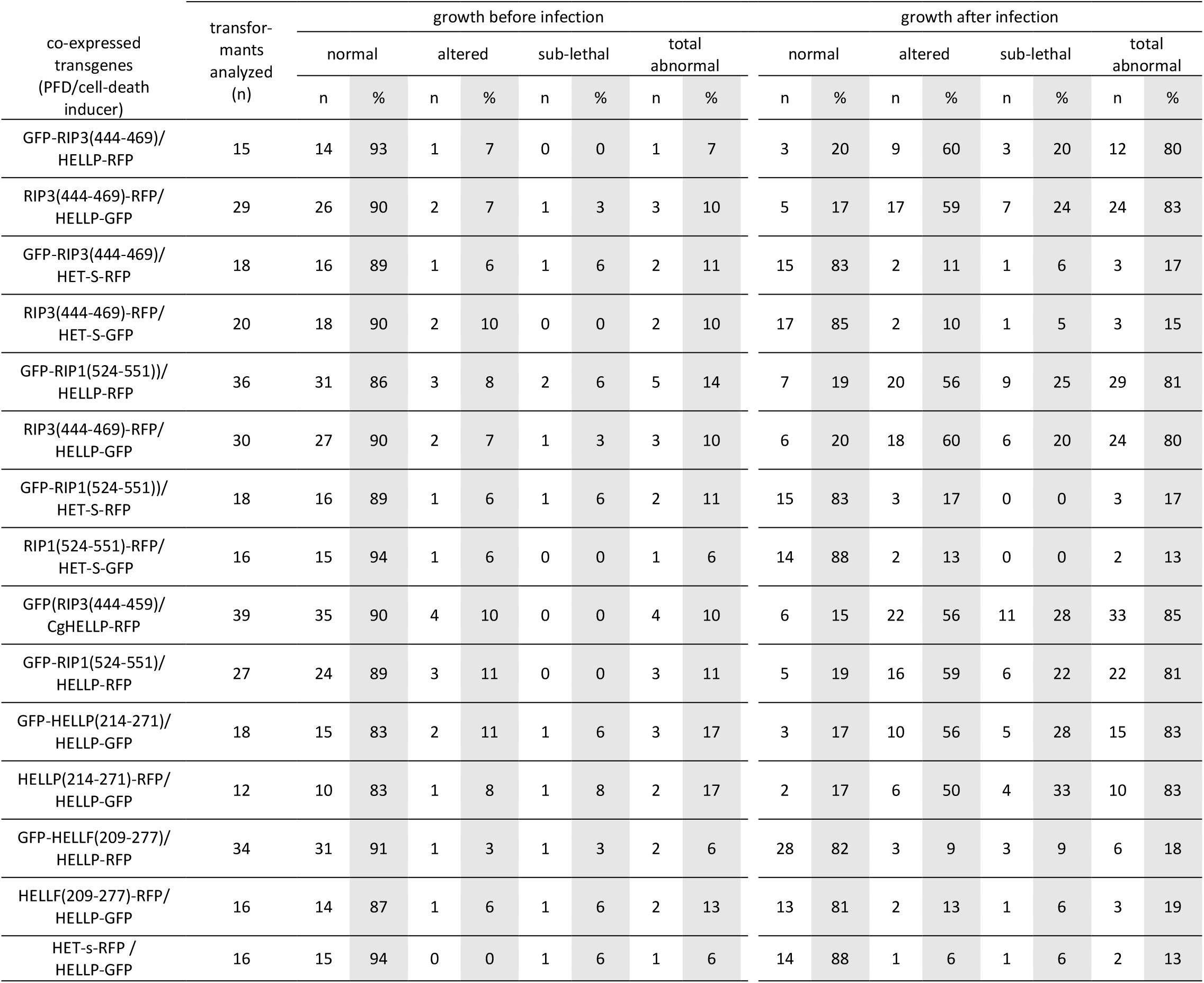
Growth alteration of strains co-expressing HELLP, CgHELLP or HET-S cell death inducing proteins and different prion forming domains prior and after prion infection.

**Fig 10.**
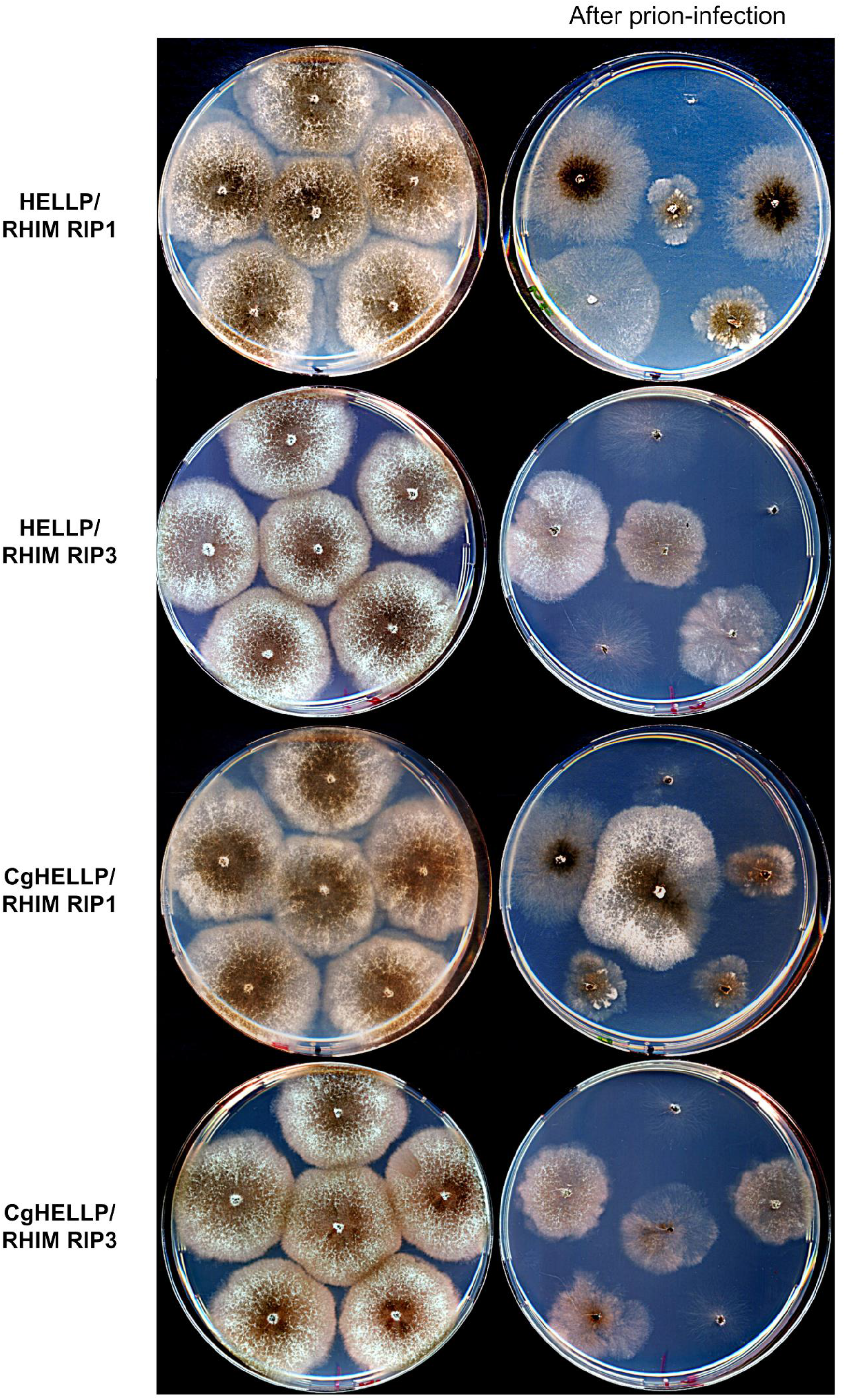
[Rhim] prions elicit self-incompatibility in strains co-expressing HELLP or CgHELLP and RIP1 or RIP3 RHIM. Comparison of growth on solid medium of strains co-expressing HELLP-RFP (HELLP) or CgHELLP-RFP (CgHELLP) and GFP-RIP1(524-551) (RIP1 RHIM) or GFP-RIP3(444-469) (RIP3 RHIM) before (left column) and after contact with a [Rhim] prion containing strain (right column, after prion-infection). Note that infection with [Rhim] prions leads to self-incompatibility with growth alterations ranging from a sublethal phenotype to a more or less stunted growth.

## Discussion

A number of innate immune pathways function following a principle of “signaling by cooperative assembly formation” (SCAF) (30). Upon recognition of PAMPs (Pathogen-associated molecular pattern) or MAMPs (Microbe-associated molecular pattern), receptors assemble into higher-order complexes or “supramolecular organizing centers” (SMOCs) (31). Following ligand recognition, the sensor modules oligomerize and recruit adaptor or effector proteins-often via homotypic interactions-that further amplify the oligomerization process and form cooperative, open-ended polymers (typically filaments). This general scheme applies to many immune signaling modules including the RIG-I-like receptors, the AIM2-like receptors and to NLRs. Formation of higher-order assemblies entitles such signaling cascade with signal amplification, sharp threshold behavior and noise reduction. In fact, this so-called prion-like polymerization process can involve nucleation of either folded globular domains such as the TIR or Death domains (32, 33) or short amyloid motifs. The first such motif is RHIM identified in the necroptosis pathway in mammals and allows for assembly of the necrosome formed by the RIP1 and RIP3 kinases (23). RHIM-like motif also regulate amyloid formation in innate immune cascades in Drosophila (24). In addition, several NLR-associated amyloid signaling motifs were identified in filamentous fungi and more recently in multicellular bacteria (18, 34). Among them is the PP-motif that shows similarity to the animal RHIM and RHIM-like motifs (22). We identify here in the species *Podospora anserina*, a PP-motif two-component system comprising a NLR termed PNT1 and a cell-death execution protein termed HELLP. As previously reported for CgHELLP from *C. globosum*, the N-terminal HELL domain of HELLP shows homology to the 4HB domain of MLKL responsible for membrane permeation in mammalian necroptosis. HELLP induced PCD thus appears to be related to a family of evolutionary widespread form of immune PCD which encompasses fungal cell death, necroptosis and cell death associated with the hypersensitive response in plants (22, 35, 36). We propose that PNT1 and HELLP constitute a third amyloid immune-related signaling cascade controlling cell fate in *P. anserina*. We show the absence of cross-induction between the three *P. anserina* systems allowing for the co-existence of three independent amyloid signaling cascades in the same species. It is believed that fungal NLRs akin to their plant and animal counterparts represent innate immune receptors that regulate response to pathogen and symbiotic interactions in fungi. The three amyloid-associated NLRs identified in Podospora share the fact that their WD or TPR repeat ligand binding domain is highly polymorphic in natural populations consistent with the proposed role as immune receptor (13, 14). Yet, except in the specific case of NLRs involved in incompatibility systems the nature of the ligands activating these NLRs are currently unknown. Compared to the two other systems, the NWD2/HET-S is characterized by the fact that this system has been exaptated as an allorecognition system in [Het-s]/HET-S incompatibility (37). It has been possible experimentally, to derive incompatibility systems analogous to [Het-s]/HET-S from both HELLP and HELLF. It will be of interest in the future to survey, natural Podospora strains to determine whether exaptation of allorecognition incompatibility might have also occurred from HELLP and HELLF. It is also of note, that the *Chaetomium globosum* and *Podospora anserina* PP-based signalosome differ in the sense that the *Chaetomium* system has three components and involves also CgSBP, a lipase also comprising a PP-motif and that is targeted to the membrane region by CgHELLP. The present study is superior to the previous study on the PP-gene cluster in the sense that HELLP function is being studied here in the native context rather than by heterologous expression (22).

So far, four mammalian proteins involved in necroptosis and five drosophila proteins also involved in immune signaling rely on RHIM-based amyloid signaling (23, 24). The fact that a range of viruses express RHIM containing proteins and that pathogenic bacteria express an enzyme that specifically cleave host RHIMs to prevent necrosome assembly highlight the crucial role of RHIM-based interactions in the host innate immune response against microbial infection (38). PP-motifs of CgHELLP and HELLP resemble RHIM motifs (Fig. 1B). This similarity is centered in the G-Φ-Q-Φ-G core of the motif. Mompeán and co-workers resolved the structure of human RIP1-RIP3 amyloid region as a serpentine fold with turns and kinks resulting in an enclosed oblong-shaped hydrophobic core stabilized by N and Q ladders and Y stacking (25). RHIM-related motif appear to represents a new class of amyloid fold distinct from the HRAM β-solenoid fold of HET-s but nonetheless presenting common features with them as with other amyloid forming domains such as conservation of G/N/Q residues. N and Q allow for formation of H-bonded N/Q ladders and G connect short adjacent β-strands (11, 39–41). The fungal PP-motifs show strong conservation of the pseudopalindromic structure centered on the Q residue along with conservation of flanking N and G residues. We find here that the RHIM motifs of RIP1 and RIP3 behave as PFDs when expressed in Podospora and thus behave analogously to PP-motifs in that respect. The simple *in vivo* Podospora model might represent a useful tool to study RHIM motif assembly and interaction for instance to screen for inducers or inhibitors of RIP1/RIP3 interactions.

Importantly, we find that [Rhim] and [π] prions cross-seed, albeit not as efficiently as RHIM motifs or fungal PP-motifs. These results suggest both a structural similarity between RHIM and PP-amyloids and substantial differences since the efficiency of prion seeding is clearly distinct from that of homotypic or intraspecific seeding. Extensive structural analyses of the amyloid fibers of HELLP or CgHELLP will be required to determine to which extend PP-based amyloid polymers share the same fold as RIP1-RIP3. It appears however that the partial cross-seeding between PP and RHIM support a model of a common evolutionary origin of the two motifs. In this context, it is interesting to note that in addition to the RHIM-like motifs described in Drosophila a RHIM-related amyloid motif termed BASS3 was also identified in multicellular bacteria (34). It might be that the RHIM-related motif represent an archetypal amyloid motif largely shared through evolution while other motifs such as HRAMs or other bacterial signaling motifs have a more restricted phylogenetic distribution.

## Materials and Methods

### Gene annotation

The Pa_5_8070 and Pa_5_8060 genes were annotated manually using two intron prediction softwares, the GENSCAN Web server at MIT (genes.mit.edu/GENSCAN.html) and the NetGene2 Web server (www.cbs.dtu.dk/services/NetGene2/), and using blastn searches against EST sequences at ncbi and RNASEQ data from (42). A single 47pb intron was identified in nucleotide position 429 to 475 of the Pa_5_8070 ORF, this newly defined ORF encodes a 271 aa protein named HELLP. A single intron of 54 pb was identified in nucleotide position 95 to 148 of the ORF of Pa_5_8060, TPR repeat number in the 3’ part of the ORF is variable in *P. anserina* strains, we described here the ORF of the S Orsay reference strain encoding a 971 aa protein named PNT1.

### Strains and plasmids

The *P. anserina* strains used in this study were wild-type *het-s* or *het-S* and the strains *Δhellp* (*ΔPa_5_8070*) *het-s°* (22) and *Δhellp (ΔPa_5_8070) Δhet-s (ΔPa_3_620) Δhellf (ΔPa_3_9900)* obtained by crossing the *Δhet-s Δhellf* strain (20) with the *Δhellp het-s°* strain. The *Δhellp het-s°* strain was used as recipient strain for the expression of molecular fusions of full length HELLP or PP motif-containing regions of HELLP and the GFP (green fluorescent protein) or RFP (red fluorescent protein). These fusions were expressed from plasmids based on the pGEM-T backbone (Promega) named pOP plasmids (22), containing either the GFP or RFP, or in a derivative of the pAN52.1 GFP vector (27), named pGB6-GFP plasmid. In both cases, the molecular fusions were under the control of the strong constitutive *P. anserina* gpd (glyceraldehyde-3-phosphate dehydrogenase) promoter. The *Δhellp het-s°* strain was transformed as described (43) with a fusion construct along with a second vector carrying a phleomycin-resistance gene ble, pPaBle (using a 10:1 molar ratio). Phleomycin-resistant transformants were selected, grown for 30 h at 26°C and screened for the expression of the transgenes using fluorescence microscopy. The *hellp* gene and the *hellp(171-271)* and *hellp(214-271)* gene fragments were amplified with the respective 5’ forward oligonucleotides 5’ ggcttaattaaATGGATCCTCTCAGTATCACAGC 3’, 5’ ggcttaattaaATGATCACAGCGACAAACGATCAG 3’ and 5’ ggcttaattaaATGAAAGTACTGCATGAATCGCGC 3’ and a same 3’ reverse oligonucleotide 5’ ggcagatcttgctccCCCCCTTCGGCCAAATGTAG 3’ (capital letters correspond to *P. anserina* genomic DNA sequences). The PCR products were cloned upstream of the GFP or RFP coding sequence in the pOP plasmids using *Pac*I/*Bgl*II restriction enzymes to generate the pOPhellp-GFP, pOPhellp-RFP, pOPhellp(171-271)-GFP, pOPhellp(171-271)-RFP, pOPhellp(214-271)-GFP and pOPhellp(214-271)-RFP vectors in which in addition to the *Bgl*II site, a two amino acid linker (GA) was introduced between the sequences encoding HELLP and GFP or RFP. *hellp(171-271)* and *hellp(214-271)* were also amplified with the respective 5’ forward oligonucleotides 5’ ggcgcgcggccgcATCACAGCGACAAACGATCAG 3’, 5’ ggcgcgcggccgcATCACAGCGACAAACGATCAG 3’ and the same 3’ reverse oligonucleotide 5’ ggcggatccCTACCCCCTTCGGCCAAATG 3’ and cloned downstream of the GFP using *Not*I/*Bam*HI restriction enzymes to generate the plasmids pGB6-GFP-hellp(171-271) and pGB6-GFP-hellp(214-271).

To investigate the co-localization of HELLP with CgHELLP or of HELLP with HELLF or HET-s, the *Δhellp Δhet-s Δhellf* strain was transformed both with vectors expressing fluorescently tagged versions of full-length or truncated HELLP described above and with fluorescently tagged versions of full-length or truncated HELLF, HET-s, HET-S or CgHELLP previously described (20, 22, 27): pOPhellf-GFP, pOPhellf(209-277)-RFP, pOPhellf(209-277)-GFP, pOPhellf(L52K)-GFP, pOPhet-s-RFP, pGPD-het-S-GFP, pOPhet-S-RFP, pGB6-GFP-Cghellp(215-278).

In the same way the sequence encoding the first 31 amino acids of *pnt1 (Pa_5_8060)* was amplified with oligonucleotides 5’ ggcttaattaaATGTCAGACAGTTATCGTTTCGGC 3’ and 5’ ggcagatcttgctccTGGCGCCTGCAGGAAAGTATTG 3’ and cloned in the pOP plasmids using *Pac*I/*Bgl*II restriction enzymes to generate the pOPpnt1(1-31)-GFP and the pOPpnt1(1-31)-RFP. The *Δhellp Δhet-s Δhellf* strain was transformed with these vectors (and pPaBle) either alone or with pOPhellp(214-271)-RFP or pGB6-GFP-hellp(214-271).

For heterologous expression in *E. coli*, we cloned *hellp(214-271)* in pET24 (Novagen) using the *Nde*I/*Xho*I restrictions sites. The gene fragment *hellp(214-271)* was amplified with oligonucleotides 5’ agccatatgAAAGTACTGCATGAATCGCGC 3’ and 5’ agcctcgagCCCCCTTCGGCCAAATGTAG 3’ and cloned in front of a polyhistidine-tag to generate pET24-hellp(214-271)-6xhis plasmid.

For the expression of RIP3(444-469) and RIP1(524-551), PCR products were amplified with 5’ ggcttaattaaATGGTTACCGGTCGTCCGCTG 3’ and 5’ ggcagatcttgctccCTGCATGGTCAGGTAGTTGTTG 3’ or 5’ ggcgcggccgcGTTACCGGTCGTCCGCTGG 3’ and 5’ ggcGGATCCTTACTGCATGGTCAGGTAGTTG 3’ for RIP3 and with 5’ ggcttaattaaATGACTGACGAATCCATCAAATACACC 3’ and 5’ ggcagatcttgctccGCCGCCGATTTCCATGTAGTT 3’ or 5’ ggcgcggccgcACTGACGAATCCATCAAATACACC 3’ and 5’ ggcGGATCCTTAGCCGCCGATTTCCATGTAGTT 3’ for RIP1 and cloned as described above in the pOP plasmids using *Pac*I/*Bgl*II restriction enzymes, or in pGB6-GFP plasmid using *Not*I/*Bam*HI restriction enzymes.

### Microscopy

*P. anserina* hyphae were inoculated on solid medium and cultivated for 24 to 72 h at 26°C. The medium was then cut out, placed on a glass slide and examined with a Leica DMRXA microscope equipped with a Micromax CCD (Princeton Instruments) controlled by the Metamorph 5.06 software (Roper Scientific). The microscope was fitted with a Leica PL APO 100X immersion lens. To observe cell-death reactions and HELLP re-localization, HELLP-GFP or HELLP-RFP expressing strains were inoculated at a distance of 2 cm from a strain expressing one of the fusion protein containing either a HELLP PFD or PNT1(1-31), RIP3(444-469) or RIP1(524-551) fused to GFP or RFP. The confrontation zone between the two strains was observed 12 to 48 hours after contact. For methylene blue staining an aqueous 0.5% solution was put directly on the mycelium for 1 min followed by washing with distilled water before observation as described above.

For HELLP fibrils observations, negative staining was performed: aggregated proteins were adsorbed onto Formvar-coated copper grids (400 mesh) and allowed to dry for 15 min in air, grids were then negatively stained 1 min with 10 μL of freshly prepared 2% uranyl acetate in water, dried with filter paper, and examined with a Hitachi H7650 transmission electron microscope (Hitachi, Krefeld, Germany) at an accelerating voltage of 120 kV. TEM was performed at the Pôle Imagerie Électronique of the Bordeaux Imaging Center using a Gatan USC1000 2k × 2k camera.

### Incompatibility assays (barrage tests)

Methods for determination of incompatibility phenotypes were previously described (20, 44). In brief, incompatibility phenotypes were determined by confronting strains on solid corn-meal agar medium and a ‘barrage’ reaction (abnormal contact lines forming upon confrontation of incompatible strains) was assessed 3 days post-contact. The [π] phenotype (acquisition of the [π] prion) was assessed as the ability of a strain to form a barrage with a wild-type strain or with a *Δhellp* bearing a transgene encoding full-length HELLP in fusion with GFP or RFP, named tester strain. Cross reaction between the different prion systems was also assessed in barrage tests as the ability of a prion containing strain (either [π], [Rhim], [Het-s] or [Φ]) to form a barrage with a strain expressing one of the full length HeLo or HELL containing proteins HELLP, CgHELLP, HELLF or HET-S. In these experiments strains previously described by Daskalov and colleagues were used (20, 22).

### Prion propagation

Methods for determination of prion formation and propagation were previously described (20, 44). Prion formation and propagation can be observed either using microscopy by monitoring the apparition of dots or using barrage tests by observing the formation of a barrage with the tester strain.

Spontaneous prion formation is first monitored as the rate of spontaneous acquired prion phenotype (dots) in the initially prion-free subculture after 5, 11 and 19 days of growth at 26°C on corn-meal agar using microscopy as described. Then transformants were re-observed at least 60 days after transformation to confirm prion acquisition. For [π] strains, barrage tests were realized 5, 19 and after at least 60 days and a strict correlation between presence of dots and barrage formation with HELLP expressing strains was always observed.

Prion formation can also be measured as the ability to propagate prions from a donor strain (containing prion) to a prion-free strain (induced strain). In practice, prion-free strains are confronted on solid corn-meal agar medium for 4 to 6 days (this step is mentioned as previous contact with in Tables) before being subcultured and observed by fluorescence microscopy and analyzed in barrage tests. As indicated in figure legends, at least 12 different transformants were used and the tests were realized in triplicates. Again for [π] strains, barrage and dots presence are strictly correlated. Prion acquisition through induction can also be visualized directly on solid medium by positioning the inocula of donor and prion-free strains closer than the tester strain (see Fig. S2) to allow prion propagation before contact with the tester strain.

It is to note that transformants were randomly tested for prion formation allowing various expression levels of the transgene (high levels of expression are usually associated with very rapid spontaneous prion formation) except for the cross conversion test (see Fig. 8 and S8) where transformants expressing moderate level of transgene were preferred to limit the rate of spontaneous transition within the timing of the experiment that could mask the prion induction. For this experiment, a statistical analysis was performed using two tails Fisher’s test to determine p-values and validate test result as indicated in the figure legend.

### Protein preparation and fibril formation

HELLP(214-271) protein was expressed in *E. coli* BL21-CodonPlus^®^-RP competent cells as insoluble proteins and purified under denaturing conditions using its terminal 6 histidine tag as previously described (45). Briefly, cells were grown at 37°C in DYT medium to 0.6 OD_600_ and expression was induced with 1 mM isopropyl β-D-1-thiogalactopyranoside. After, 4 h, cells were harvested by centrifugation, frozen at −80°C sonicated on ice in a lysis buffer (Tris 50 mM, 150 mM NaCl, pH 8) and centrifuged for 20 min at 20,000 g to remove *E. coli* contaminants. The pellet containing HELLP(214-271) in inclusion bodies was washed in the same buffer and resuspended in denaturing buffer (8M guanidinium HCl, 150 mM NaCl, and 100 mM Tris-HCl, pH 8) until complete solubilization. The lysate was incubated with Talon Resin (CLONTECH) for 1 h at 20°C, and the resin was extensively washed with 8 M urea, 150 mM NaCl, and 100 mM Tris-HCl, pH 8. The protein was eluted from the resin in the same buffer containing 200 mM imidazole. The protein HELLP(214-271) was pure as judged by sodium-dodecyl-sulfate polyacrylamide-gel electrophoreses (SDS-PAGE) followed by Coomassie-Blue staining and yield was in the range of ~2-4 mg of protein per liter of culture. To eliminate urea, elution buffer was replaced by overnight dialysis at 4°C against Milli-Q water. Fibrils formation resulted spontaneously from dialysis process followed by sample storage at 4°C for 7 days. Other conditions were used for sample preparation to test the influence of salt or acidic buffer on the fibrils formation: presence or absence of 500 mM NaCl in the dialysis buffer, replacement of dialysis by 50 times dilution in Milli-Q water or in ammonium acetate buffer 100 mM pH 4.5, they all resulted in spontaneous appearance of fibrils.

### Bioinformatics methods

Blast analyses on the *P. anserina* genome were achieved on the *Podospora anserina* Genome Project site (podospora.i2bc.paris-saclay.fr/). Sequence alignments were performed with Clustal Omega or MAFFT at (www.ebi.ac.uk) and edited with Jalview (www.jalview.org/). To generate the HMM profile signatures for PP and RHIM, psi-blast searches were performed at (blast.ncbi.nlm.nih.gov) using the HELLP PP sequence and the RHIM motif of human RIP3 and matching sequences were aligned with Clustal Omega and the alignment was analyzed with Skylign (skylign.org). Secondary structure predictions were performed with PSIPRED (bioinf.cs.ucl.ac.uk/psipred/). Hidden Markov Model searches were performed using HHPred (toolkit.tuebingen.mpg.de) and Jackhhmer (www.ebi.ac.uk/Tools/hmmer/search/phmmer), both with default settings. The prediction of transmembrane helix was performed with the TMHMM server (www.cbs.dtu.dk/services/TMHMM).

## Author Contributions

**Conceptualization:** Thierry Bardin, Asen Daskalov, Sven J. Saupe, Virginie Coustou

**Data curation:** Thierry Bardin, Asen Daskalov, Sophie Barrouilhet, Sven J. Saupe, Virginie Coustou

**Formal analysis:** Thierry Bardin, Asen Daskalov, Sophie Barrouilhet, Sven J. Saupe, Virginie Coustou

**Funding acquisition:** Sven J. Saupe

**Investigation:** Thierry Bardin, Asen Daskalov, Sophie Barrouilhet, Alexandra Granger-Farbos, Bénédicte Salin, Corinne Blancard, Virginie Coustou

**Methodology:** Thierry Bardin, Asen Daskalov, Alexandra Granger-Farbos, Bénédicte Salin, Corinne Blancard, Sven J. Saupe, Virginie Coustou

**Project administration:** Sven J. Saupe, Virginie Coustou

**Resources:** Alexandra Granger-Farbos, Bénédicte Salin, Corinne Blancard, Sven J. Saupe, Virginie Coustou

**Supervision:** Sven J. Saupe, Virginie Coustou

**Validation:** Thierry Bardin, Asen Daskalov, Sven J. Saupe, Virginie Coustou

**Visualization:** Thierry Bardin, Asen Daskalov, Sven J. Saupe, Virginie Coustou

**Writing:** Sven J. Saupe, Virginie Coustou

## Supporting information

### Tables

**Table S1.** [π] prion propagation assayed in barrage tests.

| tested strain<br>(prion recipient) |  | previous contact with<br>(prion donor) |  | tester strains |  |  |
| --- | --- | --- | --- | --- | --- | --- |
| recipient strain | transgene | recipient strain | transgene | *$\Delta hellp \Delta het-s \Delta hellf$* |  | recipient strain |
|  |  |  |  | HELLP-GFP | HELLP-RFP | transgene |
| *$\Delta hellp$*<br>**het-s*<sup>°</sup>* | GFP-HELLP<br>(214-271)<br>[$\pi^*$] | *$\Delta hellp$*<br>**het-s*<sup>°</sup>* | no transgene | 0/12 | 0/12 | |
| | | | GFP-HELLP(214-271) [$\pi$] | 12/12 | 12/12 | |
| | | | HELLP (214-271)-GFP [$\pi$] | 12/12 | 12/12 | |
| | | | HELLP (214-271)-RFP [$\pi^*$] | 0/12 | 0/12 | |
| | | | HELLP (214-271)-RFP [$\pi$] | 12/12 | 12/12 | |
| | | | HELLP (171-271)-GFP [$\pi$] | 12/12 | 12/12 | |
| | | | GFP-HELLP (171-271) [$\pi$] | 12/12 | 12/12 | |

The table gives the number of transformants producing a barrage reaction (after contact with the given prion donor strain), to two different tester strains expressing full-length HELLP (either as GFP or RFP fusion). For each transgene, 12 different transformants were tested and the experiment done in triplicate. All triplicates were consistent.

**Table S2.**
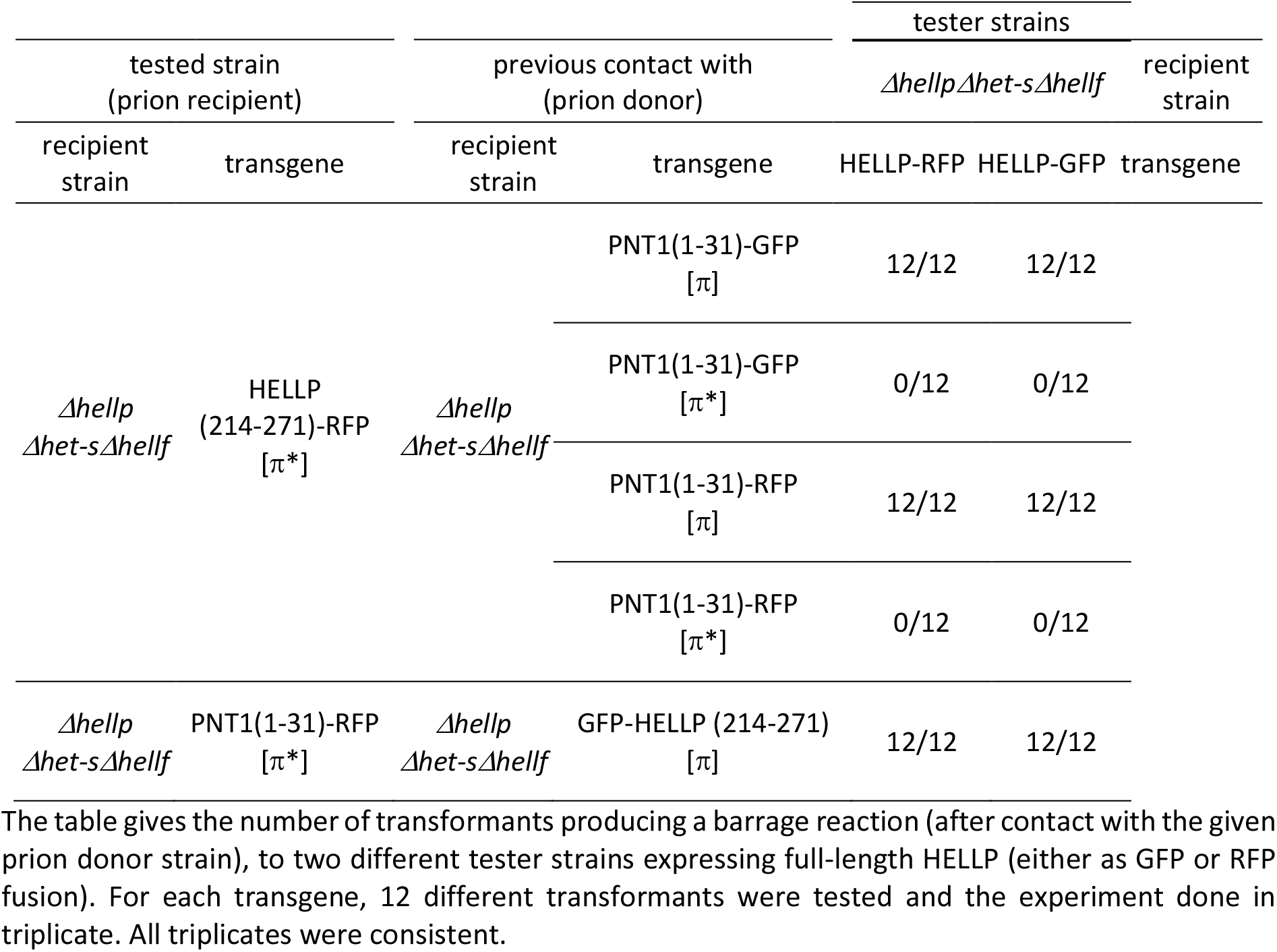
Test of cross-conversion between [π]^PNT1(1-31)^ and [π]^HELLP(214-271)^ prions.

**Table S3.** Assay in barrage tests of [π*] conversion by [Het-s] and [Φ] prions.

| tested strain<br>(prion recipient) |  | previous contact with<br>(prion donor) |  | tester strains |  | recipient strain |
| --- | --- | --- | --- | --- | --- | --- |
| recipient strain | transgene | recipient strain | transgene | HELLP-RFP | HELLP-GFP | transgene |
| *$\Delta hellp$*<br>*$\Delta het-s \Delta hellf$* | HELLP<br>(214-271)-GFP<br>[$\pi^*$] | *$\Delta hellp$*<br>*$\Delta het-s \Delta hellf$* | HET-s-RFP<br>[Het-s] | 0/12 | 0/12 | |
| | | | GFP-HELLF(209-277)<br>[$\Phi$] | 0/12 | 0/12 | |
| | | | GFP-HELLP (214-271)<br>[$\pi$] | 12/12 | 12/12 | |
| *$\Delta hellp$*<br>*$\Delta het-s \Delta hellf$* | HELLP<br>(214-271)-RFP<br>[$\pi^*$] | *$\Delta hellp$*<br>*$\Delta het-s \Delta hellf$* | HET-s-RFP<br>[Het-s] | 0/12 | 0/12 | |
| | | | GFP-HELLF(209-277)<br>[$\Phi$] | 0/12 | 0/12 | |
| | | | GFP-HELLP (214-271)<br>[$\pi$] | 12/12 | 12/12 | |

**Table S4.** Test of cross-induction of incompatibility between three *P. anserina* amyloid signaling systems.

| tested strains |  | tester strain |  |  | recipient strain |
| --- | --- | --- | --- | --- | --- |
|  |  | *ΔhellpΔhetsΔhellf* |  |  |  |
| recipient strain | transgene | HET-S-GFP | HELLF-GFP | HELLP-GFP | transgene |
| *ΔhellpΔhet-sΔhellf* | HET-s-RFP | 18/18 | 0/18 | 0/18 |  |
| *Δhet-s* | HET-s(218-289)-GFP | 18/18* | 0/18 | 0/18 |  |
| *ΔhellpΔhet-sΔhellf* | HELLF(209-277)-RFP | 0/18 | 18/18 | 0/18 |  |
| *ΔhellpΔhet-sΔhellf* | GFP-HELLF(209-277) | 0/18 | 18/18 | 0/18 |  |
| *ΔhellpΔhet-sΔhellf* | HELLF(L52K)-GFP | 0/18 | 18/18 | 0/18 |  |
| *ΔhellpΔhet-sΔhellf* | HELLP(214-271)-GFP | 0/18 | 0/18 | 18/18 |  |
| *ΔhellpΔhet-sΔhellf* | GFP-HELLP(214-271) | 0/18 | 0/18 | 18/18 |  |
| *Δhellp het-s°* | GFP-CgHELLP(215-278) | 0/18 | 0/18 | 18/18 |  |

The table gives the number of transformants producing a barrage reaction to the given tester strains. For each transgene, 18 different transformants were tested and the experiment was done in triplicate. All triplicate were consistent. \*As noted previously, HET-s(218-289)-GFP produces an attenuated barrage reaction to HET-S.

**Table S5.**
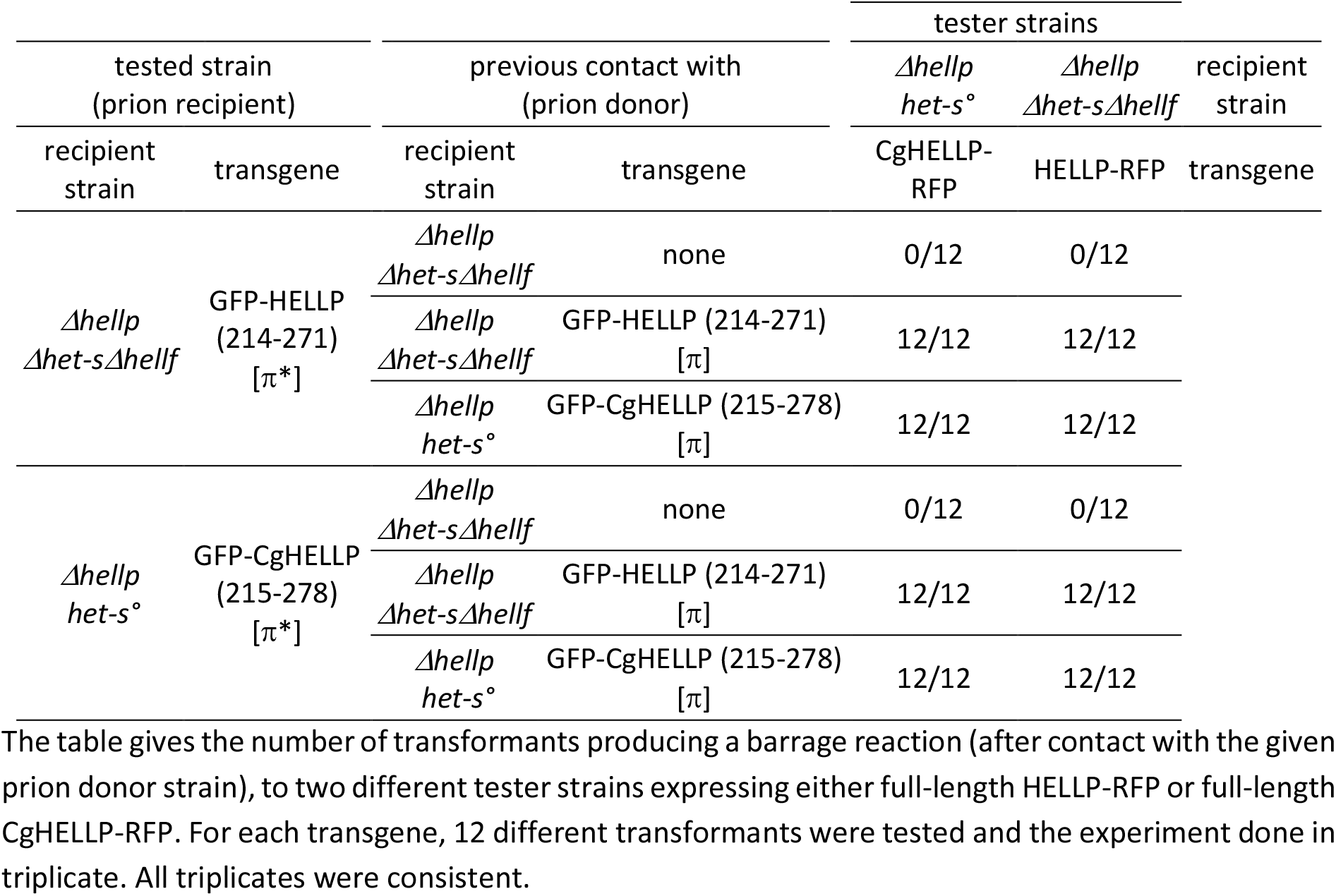
Test of cross-conversion between [π]^Pa^ and [π]^Cg^ prions and induction HELLP and CgHELLP toxicity.

**Table S6.** Cross-induction of HELLP and CgHELLP cell death activity by [π] prions.

| tested strains |  | tester strains |  |  |  |  | recipient strain |
| --- | --- | --- | --- | --- | --- | --- | --- |
|  |  | *Δhellp*<br>*Δhet-sΔhellf* | *Δhellp*<br>*Δhet-sΔhellf* | *Δhellp*<br>*Δhet-sΔhellf* | wt | *Δhellp*<br>*het-s°* |  |
| Recipient strain | Transgene | none | HELLP-RFP | HELLP-GFP | none | CgHELLP-RFP | transgene |
| *Δhellp*<br>*Δhet-sΔhellf* | none | 0/18 | 0/18 | 0/18 | 0/18 | 0/18 |  |
| *Δhellp*<br>*Δhet-sΔhellf* | GFP-HELLP (214-271) | 0/18 | 18/18 | 18/18 | 18/18<sup>a</sup> | 18/18 |  |
| *Δhellp*<br>*Δhet-sΔhellf* | GFP-HELLP (171-271) | 0/18 | 18/18 | 18/18 | 18/18<sup>a</sup> | 18/18 |  |
| *Δhellp*<br>*Δhet-sΔhellf* | HELLP (214-271)-RFP | 0/18 | 18/18 | 18/18 | 18/18<sup>a</sup> | 18/18 |  |
| *Δhellp het-s°* | GFP-CgHELLP (215-278) | 0/18 | 18/18 | 18/18 | 0/18 | 18/18 |  |
| *Δhellp het-s°* | CgHELLP (170-278)-RFP | 0/18 | 18/18 | 18/18 | 0/18 | 18/18 |  |

The table gives the number of transformants producing a barrage reaction to the given tester strains (<sup>a</sup> barrage was attenuated). For each transgene, 18 different transformants were tested and the experiment was done in triplicate. All triplicates were consistent.

**Table S7.**
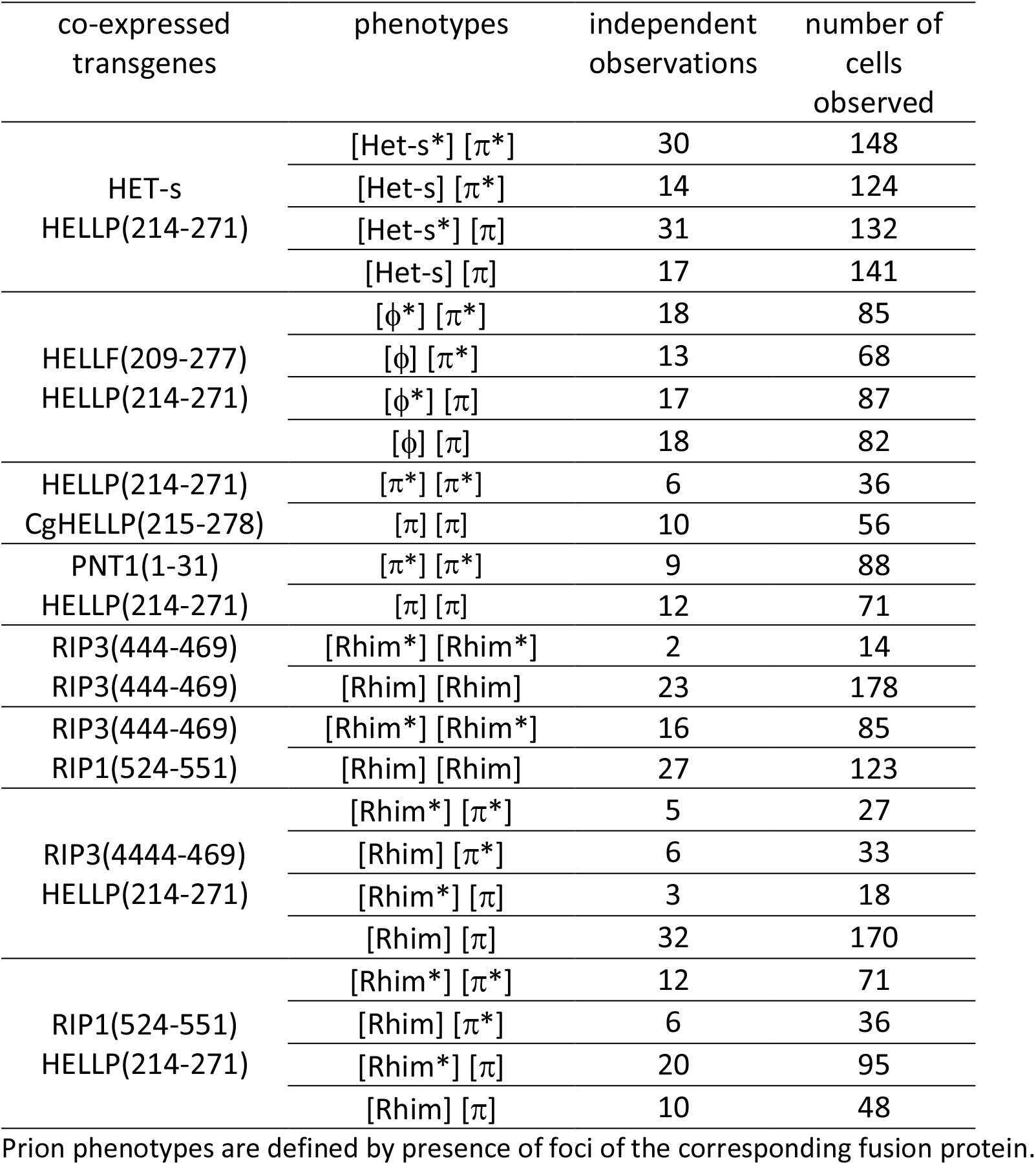
Occurrence of the different phenotypes observed using fluorescence microscopy during co-expression experiments.

**Table S8.**
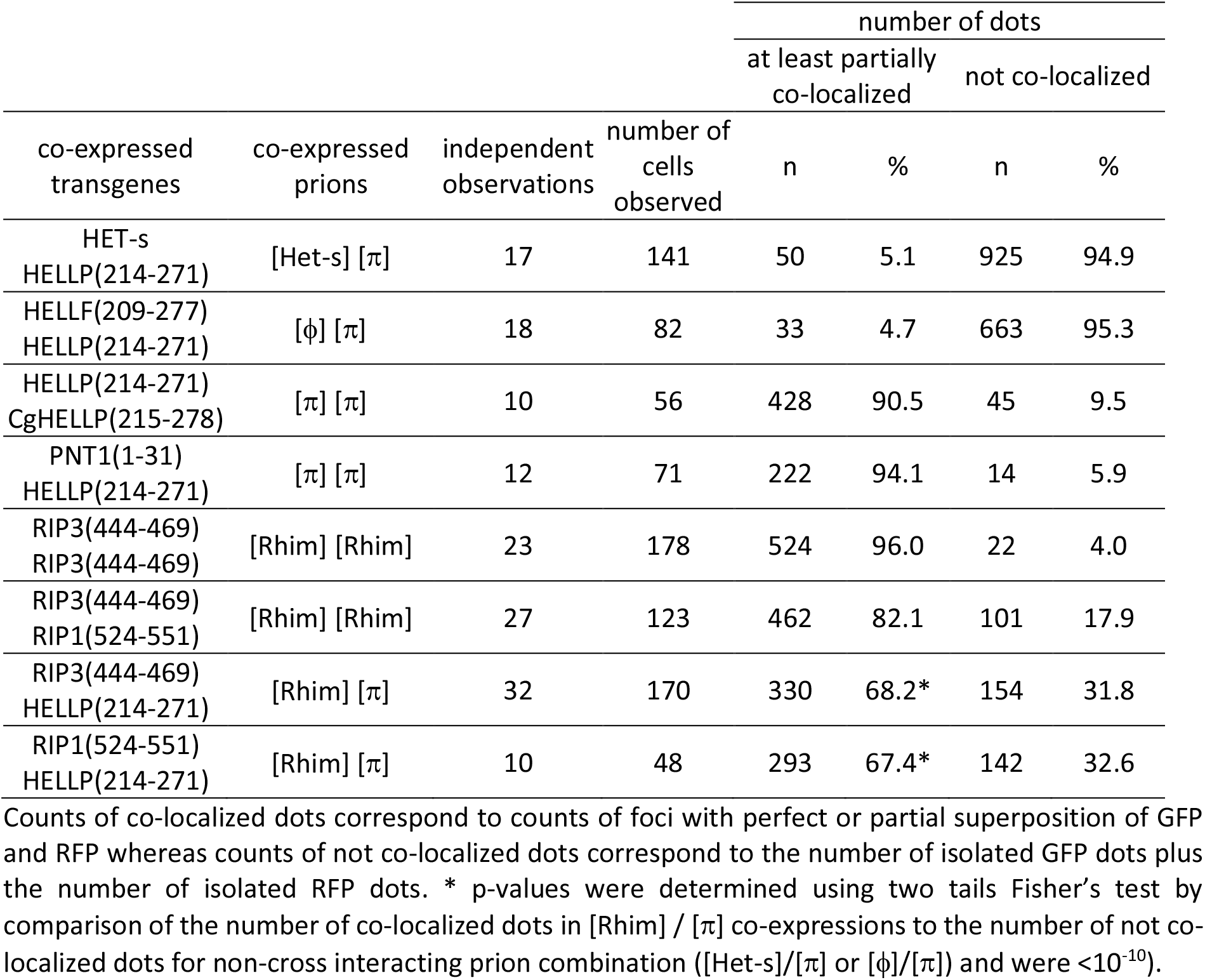
Count of co-localized foci observed by fluorescence microscopy during co-expression experiments.

| co-expressed transgenes | co-expressed prions | independent observations | number of cells observed | number of dots |  |  |  |
| --- | --- | --- | --- | --- | --- | --- | --- |
|  |  |  |  | at least partially co-localized |  | not co-localized |  |
|  |  |  |  | n | % | n | % |
| HET-s<br>HELLP(214-271) | [Het-s] [$\pi$] | 17 | 141 | 50 | 5.1 | 925 | 94.9 |
| HELLF(209-277)<br>HELLP(214-271) | [$\phi$] [$\pi$] | 18 | 82 | 33 | 4.7 | 663 | 95.3 |
| HELLP(214-271)<br>CgHELLP(215-278) | [$\pi$] [$\pi$] | 10 | 56 | 428 | 90.5 | 45 | 9.5 |
| PNT1(1-31)<br>HELLP(214-271) | [$\pi$] [$\pi$] | 12 | 71 | 222 | 94.1 | 14 | 5.9 |
| RIP3(444-469)<br>RIP3(444-469) | [Rhim] [Rhim] | 23 | 178 | 524 | 96.0 | 22 | 4.0 |
| RIP3(444-469)<br>RIP1(524-551) | [Rhim] [Rhim] | 27 | 123 | 462 | 82.1 | 101 | 17.9 |
| RIP3(444-469)<br>HELLP(214-271) | [Rhim] [$\pi$] | 32 | 170 | 330 | 68.2* | 154 | 31.8 |
| RIP1(524-551)<br>HELLP(214-271) | [Rhim] [$\pi$] | 10 | 48 | 293 | 67.4* | 142 | 32.6 |

Counts of co-localized dots correspond to counts of foci with perfect or partial superposition of GFP and RFP whereas counts of not co-localized dots correspond to the number of isolated GFP dots plus the number of isolated RFP dots. \* p-values were determined using two tails Fisher's test by comparison of the number of co-localized dots in [Rhim] / [ $\pi$ ] co-expressions to the number of not co-localized dots for non-cross interacting prion combination ([Het-s]/[ $\pi$ ] or [ $\phi$ ]/[ $\pi$ ]) and were $<10^{-10}$ ).

### Supplementary figures

**Fig S1.**
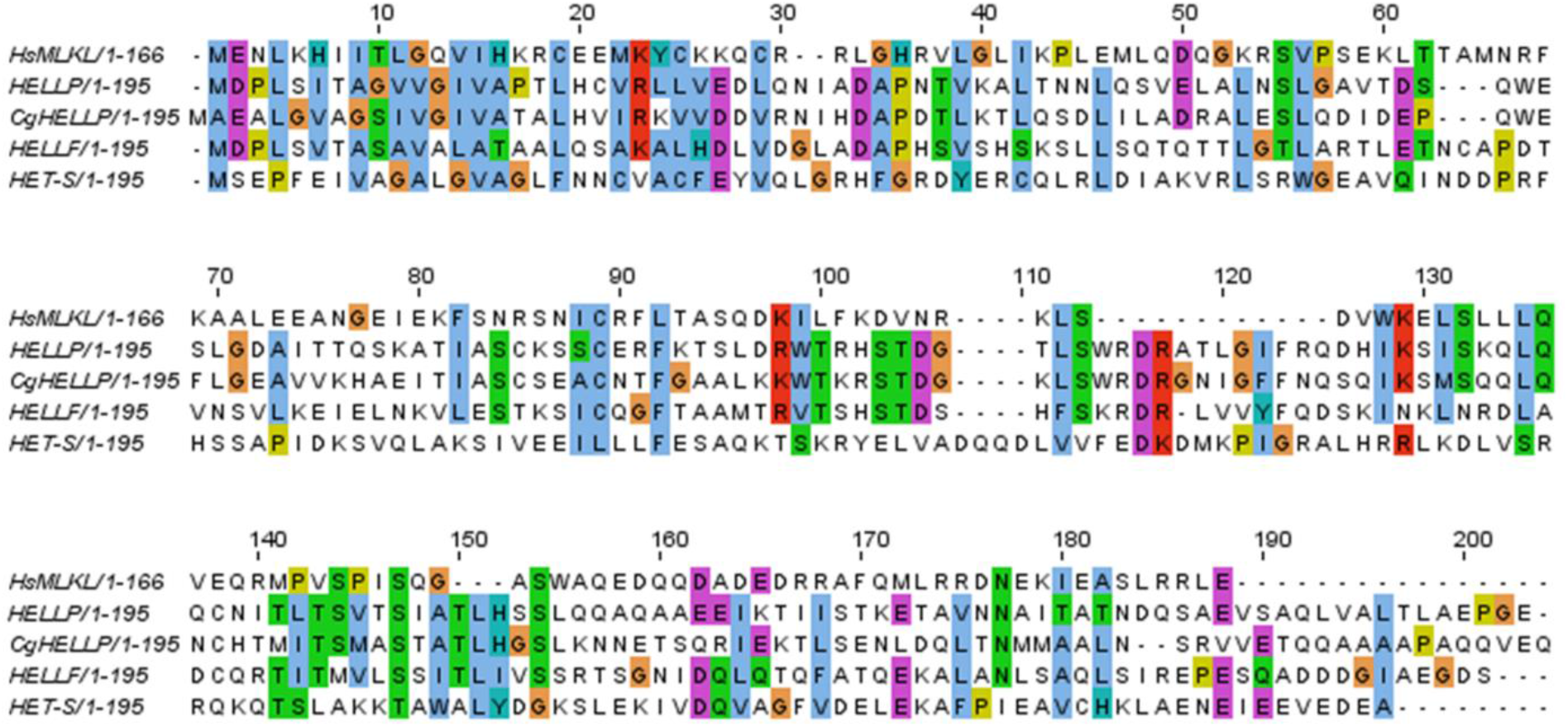
Homology between HeLo or HELL domains of *P. anserina* and 4HB domain of human MLKL. Alignment of the HET-S HeLo domain and the HELLF and HELLP HELL domains of *P. anserina* with the 4HB region of the human MLKL. Alignment was generated with clustal omega with default settings.

**Fig S2.**
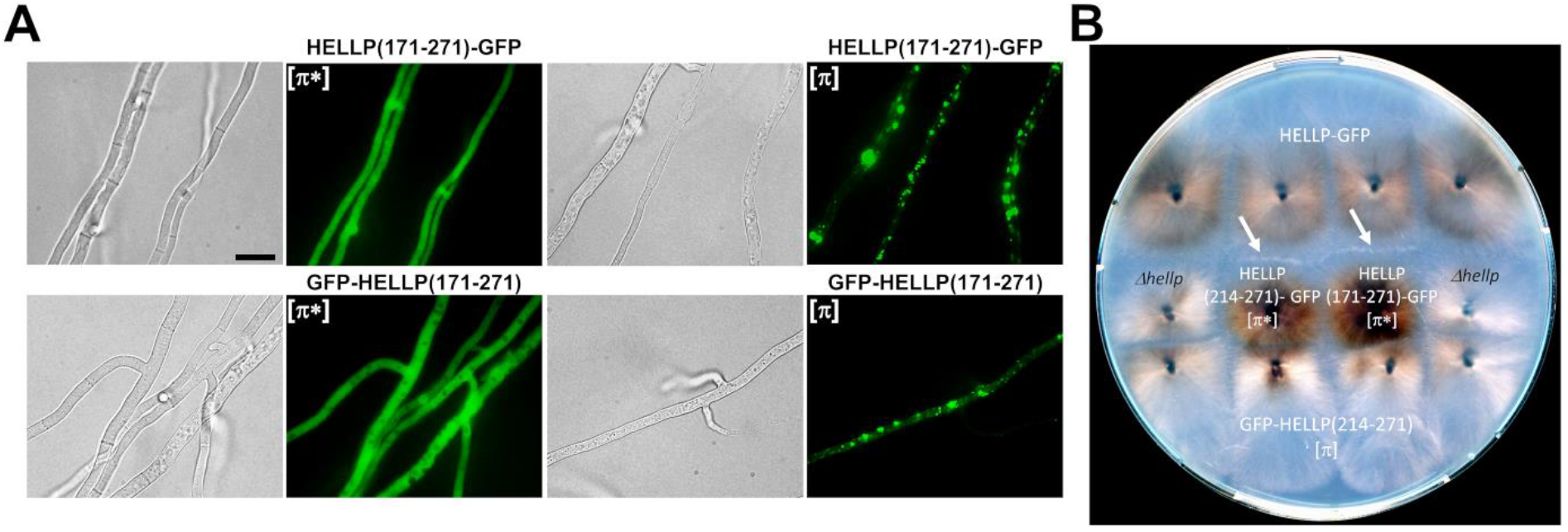
[π] prion propagation of longer HELLP constructs. (A) Micrographs of *P. anserina* strains expressing HELLP(171-271)-GFP or GFP-HELLP(171-271) as indicated above each micrograph (scale bar is 5 μm). Transformants presenting a diffuse fluorescence in the non-prion state designated [π*] (left) and dot-like fluorescence after contact with a [π] strain (right). (B) Barrage tests showing conversion of [π*] strains (expressing either HELLP(214-271)-GFP or HELLP(171-271)-GFP) to the [π] phenotype by contact with strains expressing GFP-HELLP(214-271) in the [π] state.

**Fig S3.**
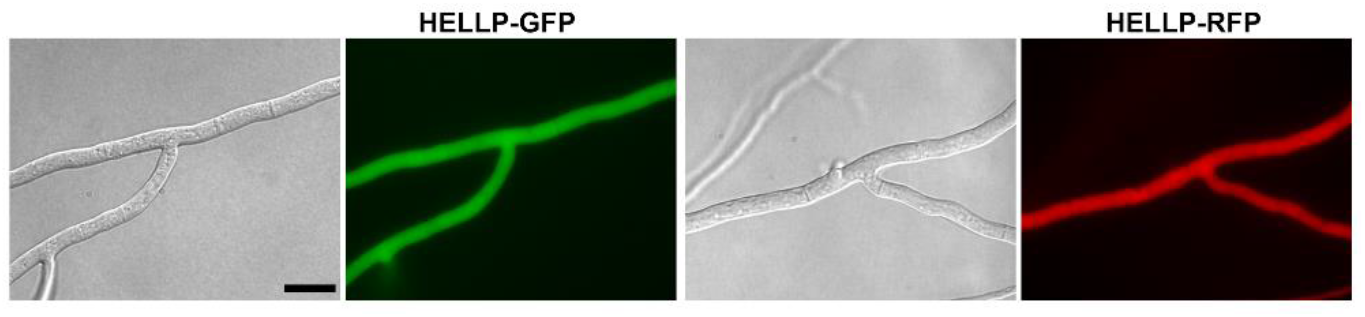
HELLP full-length diffuse and cytoplasmic localization. Micrographs of strains expressing HELLP-GFP or HELLP-RFP fusion proteins. The fluorescence signal is diffuse and cytoplasmic and remains stable, (scale bar is 5μm).

**Fig S4.**
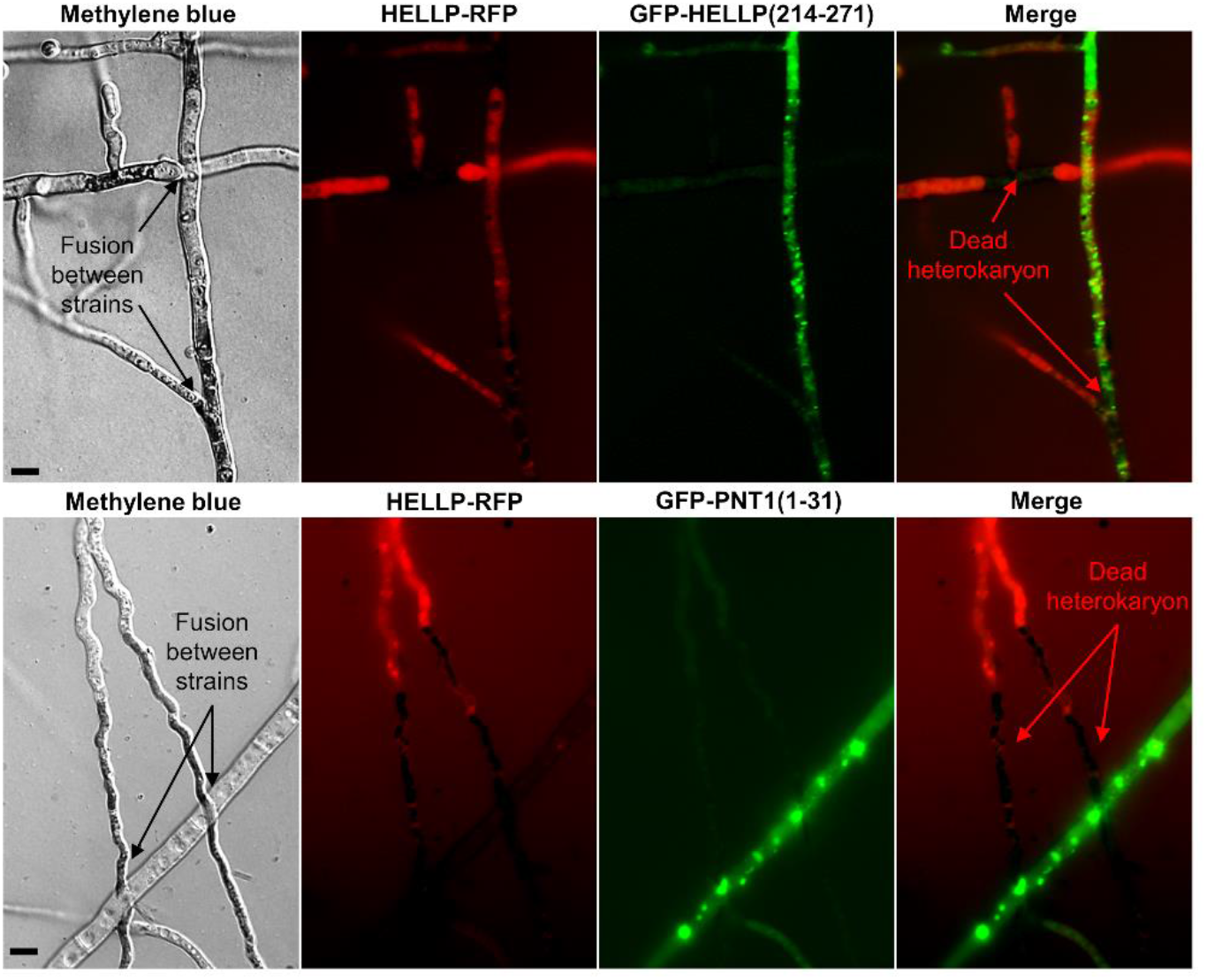
Death of fusion cells containing [π] prions and HELLP full-length proteins. Micrographs showing contact zone between HELLP-RFP and GFP-HELLP(214-271) (Top panel) or GFP-PNT1(1-31) (Bottom panel) expressing strains after Methylene blue staining. Note that heterokaryotic fusion cells are stained (appeared in black) indicating their death, (scale bar is 2 μm).

**Fig S5.**
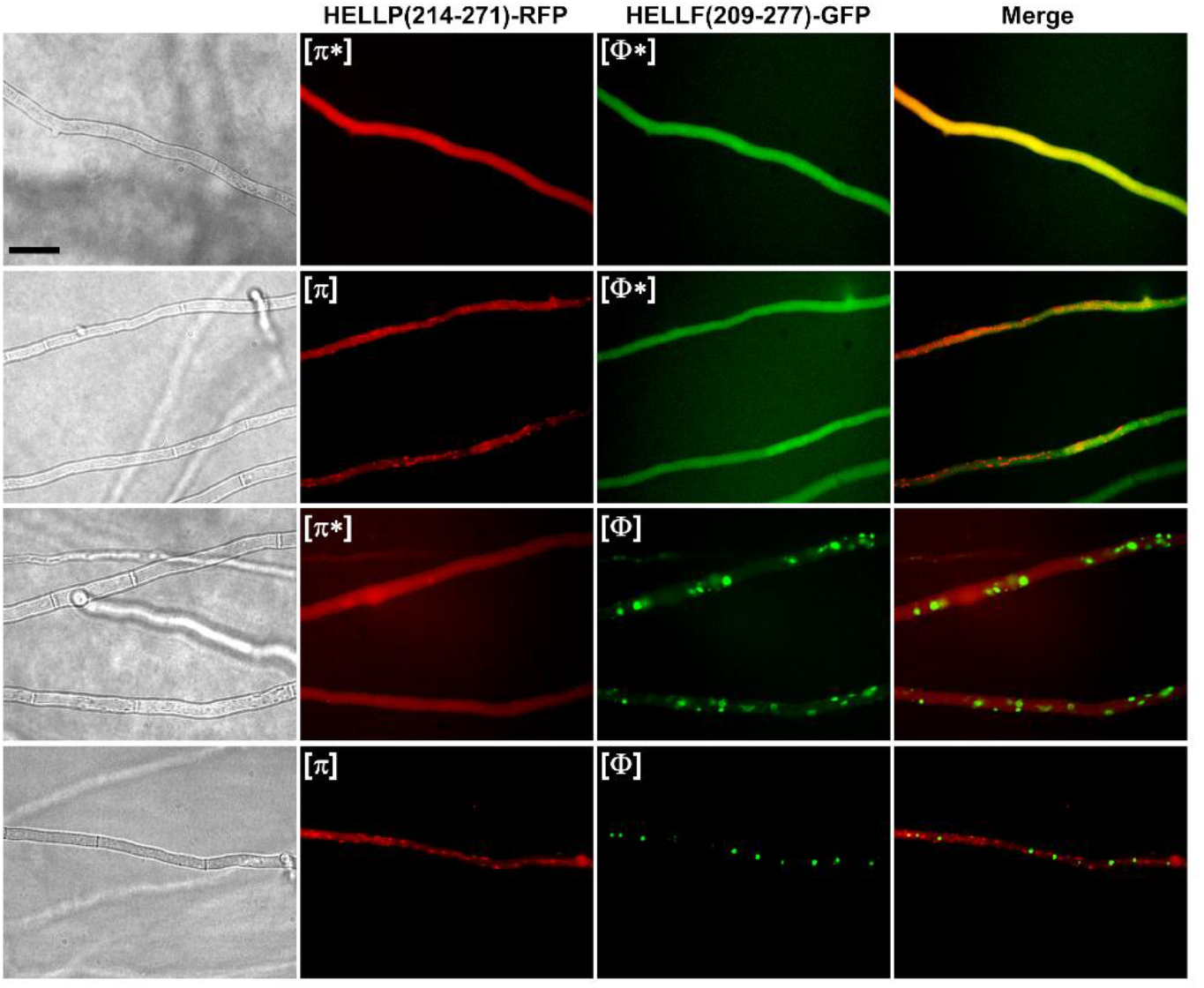
Absence of co-localization HELLP(214-271)-RFP and HELLF(209-277)-GFP foci. Micrographs of strains co-expressing HELLP(214-271)-RFP ([π*] or [π] state) and HELLF(209-277)-GFP ([Φ*] or [Φ] state). Note that HELLP(214-271)-RFP and HELLF(209-277)-GFP foci do not co-localize ([Φ] and [π]) and the existence of cells in which one of the fusion protein forms foci while the other remains diffuse, (scale bar is 5 μm).

**Fig S6.**
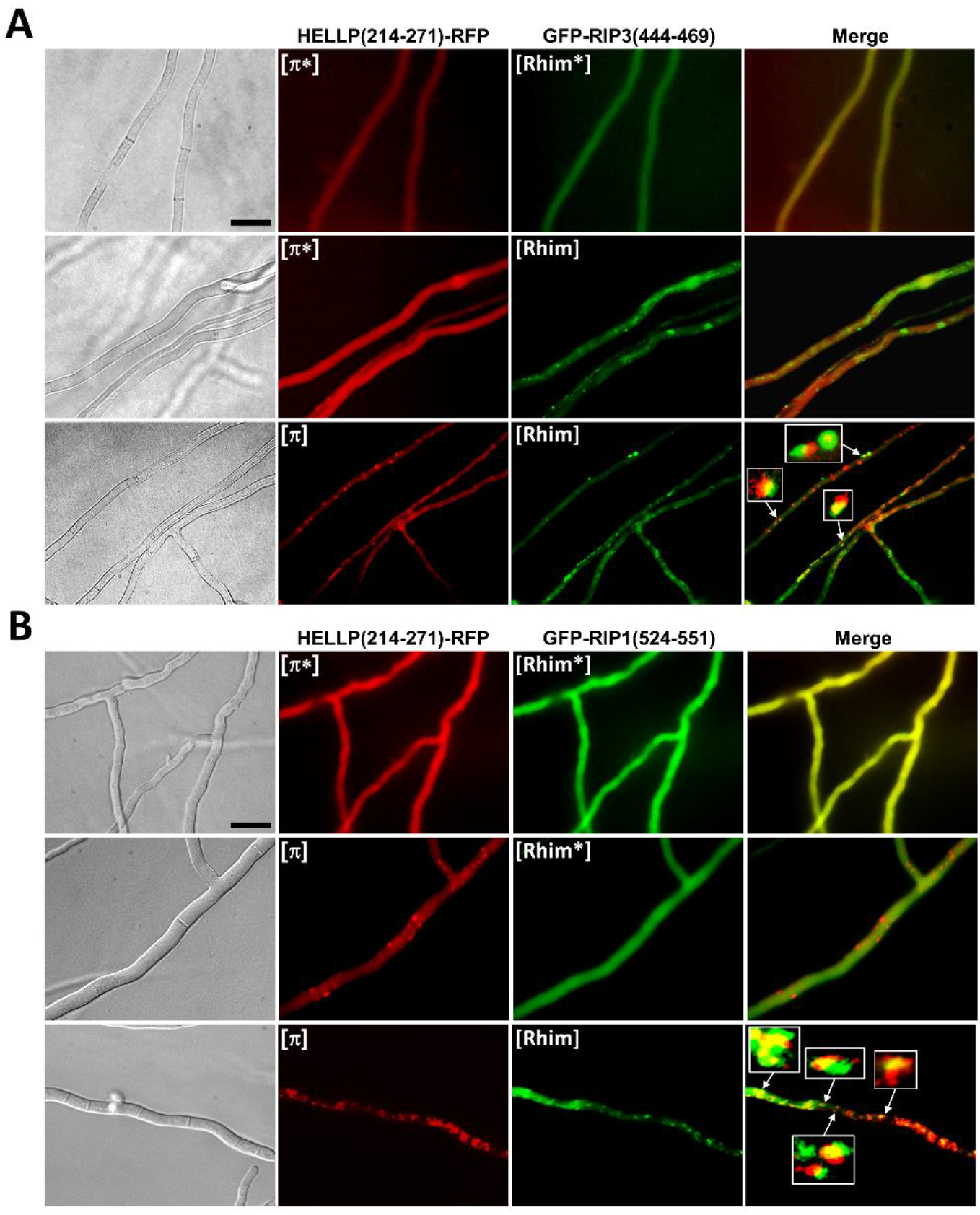
Partial co-localization HELLP(214-271)-RFP foci and GFP-RIP1(524-551) and GFP-RIP3(444-469) foci. (A) Micrographs of strains co-expressing GFP-RIP3(444-469) ([Rhim*] or [Rhim] states) and HELLP(214-271)-RFP ([π*] or [π] states). (B) Micrographs of strains co-expressing GFP-RIP1(524-551) ([Rhim*] or [Rhim] states) and HELLP(214-271)-RFP ([π*] or [π] states). In both cases, note the co-existence on the second lane of [Rhim] and [π*] (A) or [Rhim*] and [π] (B),. Note the partial co-localization of the foci; some of the foci were magnified to show incomplete overlap of the fusion protein, (scale bar is 5 μm).

**Fig S7.**
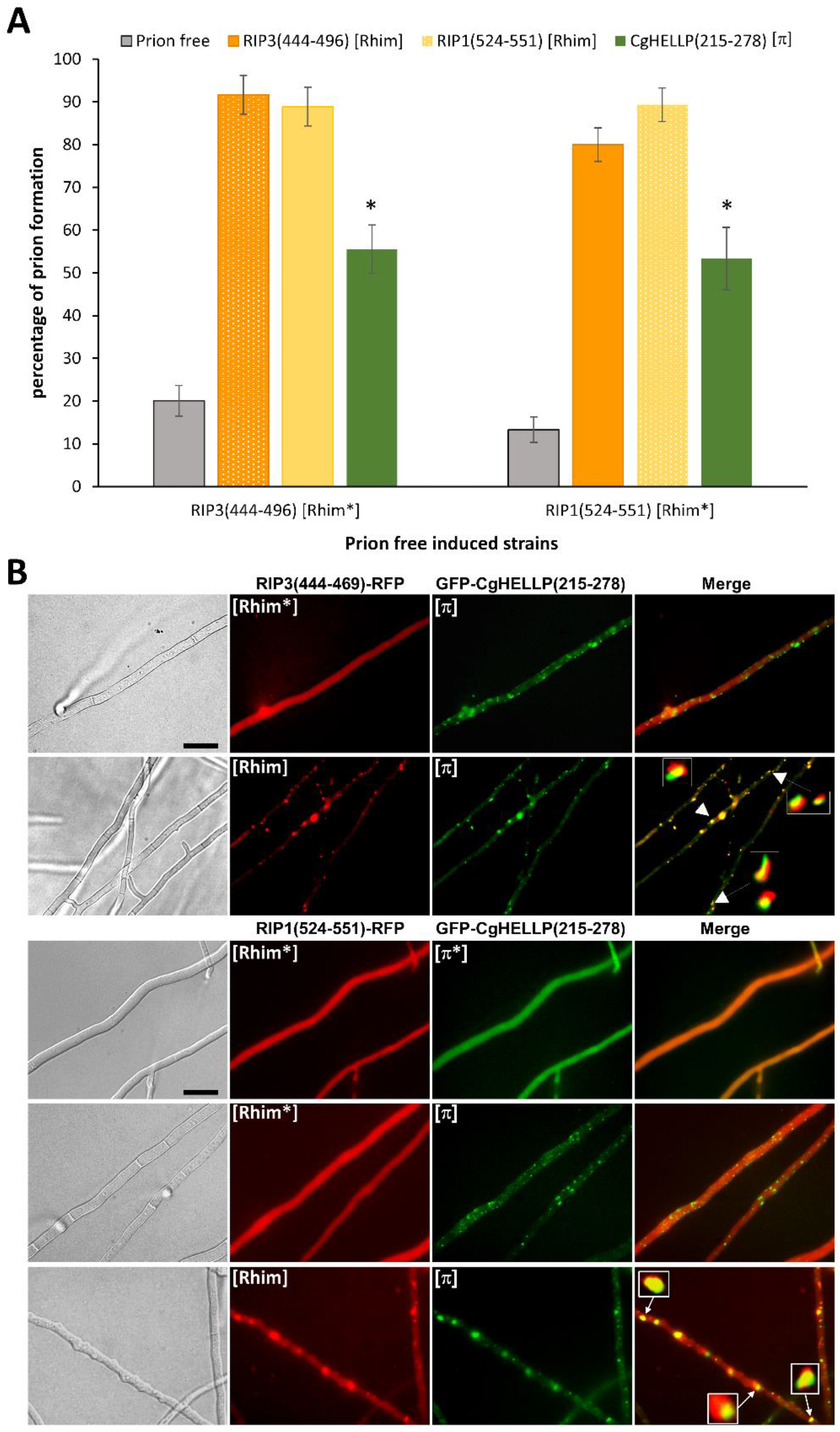
Interaction between human RIP3 or RIP1 RHIM region and the CgHELLP PFD. (A) Induction of [Rhim] prion by cross conversion of [Rhim*] by [π]^Cg^ or [Rhim] prions. Histogram representing the percentage of [Rhim*] strains displaying [Rhim] phenotype after contact with a *ΔhellpΔhet-sΔhellf* prion free strain (control), a GFP-CgHELLP(215-278) [π] containing strain or a GFP-RIP3(444-469) or GFP-RIP1(524-551) [Rhim] containing strains (indicated on top). Strains phenotype after induction was determined by monitoring the acquisition of dot-like aggregates by fluorescence microscopy. Homotypic interactions shown with striped lines. Percentages of prion formation were expressed as the mean value ± standard deviation for 3 to 5 independent experiments on 4 different transformants. P-values for cross conversion were determined using two tails Fisher’s test by comparison of the number of prion free and prion containing strains obtained after induction by the prion free control or by the heterotypic prion containing strain and are <10^−8^. (B) Partial co-localization of [π] Cg and [Rhim] prions *in vivo*. Micrographs of strains co-expressing RIP3(444-469)-RFP or RIP1(524-551)-RFP ([Rhim*] or [Rhim] states) and GFP-CgHELLP(215-278) ([π*] and [π] state). Note the co-existence of [Rhim*] and [π], this situation is only observable for a short period of time, showing a low conversion rate of [Rhim*] by [π] prion. Note the partial co-localization of the two prions, some of the dots were zoomed to show incomplete overlapping, (scale bar is 5 μm).

**Fig S9.**
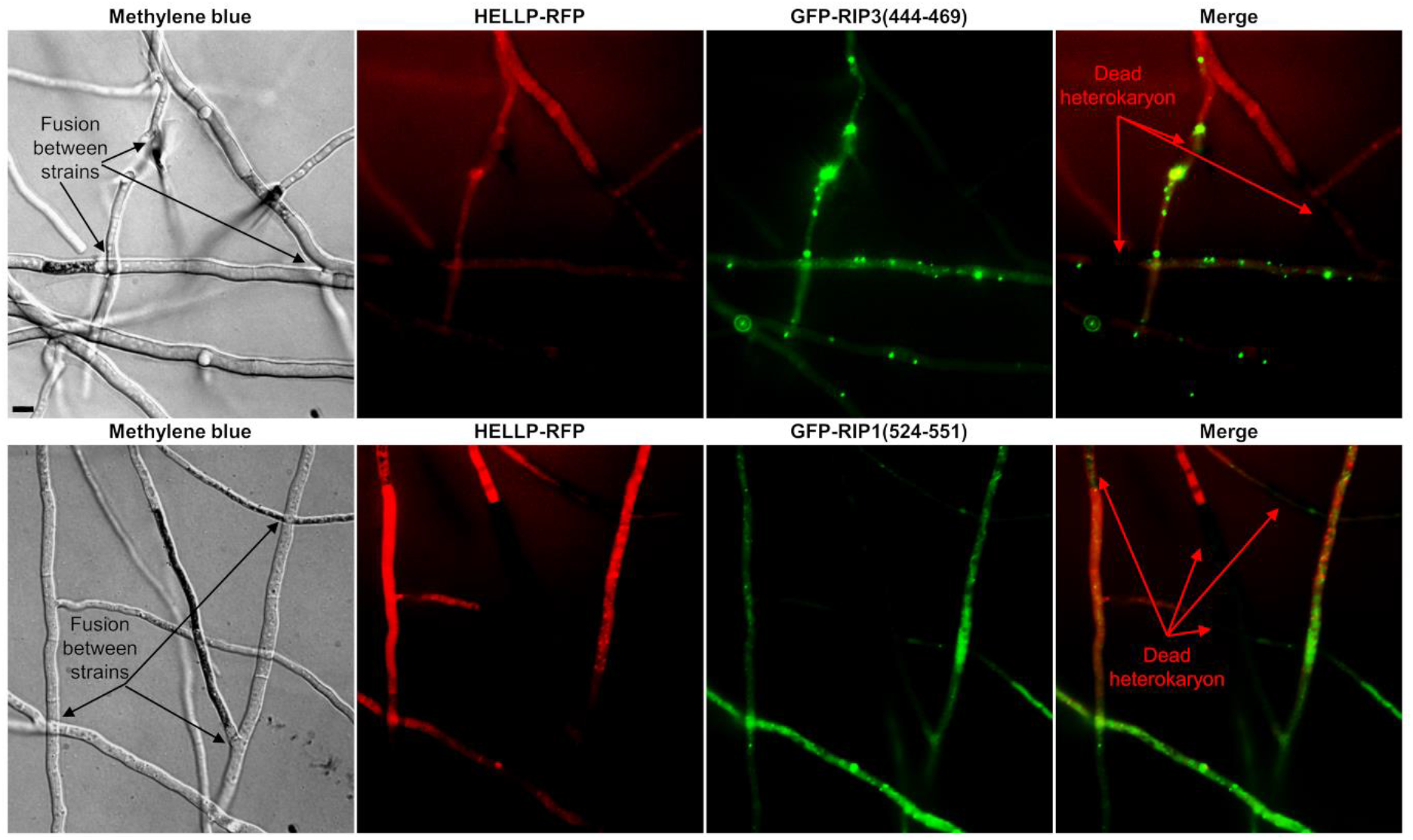
Cell death occurring in fusion cells between [Rhim] and HELLP expressing strains. Micrographs showing confrontations between HELLP-RFP and GFP-RIP3(444-469) (Upper panel) or GFP-RIP1(524-551) (lower panel) expressing strains after methylene blue staining. Note that heterokaryotic fusion cells are stained indicating their death, (scale bar is 2 μm).

## Notes

### Competing Interest Statement

The authors have declared no competing interest.

